# Chromosome-scale genome assemblies of aphids reveal extensively rearranged autosomes and long-term conservation of the X chromosome

**DOI:** 10.1101/2020.03.24.006411

**Authors:** Thomas C. Mathers, Roland H. M. Wouters, Sam T. Mugford, David Swarbreck, Cock Van Oosterhout, Saskia A. Hogenhout

## Abstract

Large-scale chromosome rearrangements are arguably the most dramatic type of mutations, often leading to rapid evolution and speciation. However, chromosome dynamics have only been studied at the sequence level in a small number of model systems. In insects, Diptera (flies and mosquitoes) and Lepidoptera (butterflies and moths) have high levels of chromosome conservation. Whether this truly reflects the diversity of insect genome evolution is questionable given that many species exhibit rapid karyotype evolution. Here, we investigate chromosome evolution in aphids – an important group of hemipteran plant pests – using newly generated chromosome-scale genome assemblies of the green peach aphid (*Myzus persicae*) and the pea aphid (*Acyrthosiphon pisum*), and a previously published chromosome-scale assembly of the corn-leaf aphid (*Rhopalosiphum maidis*). We find that aphid autosomes have undergone dramatic reorganisation over the last 30 million years, to the extent that chromosome homology cannot be determined between aphids from the tribes Macrosiphini (*M. persicae* and *A. pisum*) and Aphidini (*R. maidis*). In contrast, gene content of the aphid sex (X) chromosome remained unchanged despite rapid sequence evolution, low gene expression and high transposable element load. To test whether rapid evolution of genome structure is a hallmark of Hemiptera, we compared our aphid assemblies to chromosome-level assemblies of two blood-feeding Hemiptera (*Rhodnius prolixus* and *Triatoma rubrofasciata*). Despite being more diverged, the blood-feeding hemipterans have conserved synteny and we detect only two chromosome fusion or fission events. The exceptional rate of structural evolution of aphid autosomes renders them an important emerging model system for studying the role of large-scale genome rearrangements in evolution.

## Introduction

Mutation generates genomic novelty upon which natural selection and genetic drift can act to drive evolutionary change (Charlesworth 2009; Lynch *et al.* 2016; Charlesworth and Charlesworth 2017; Good *et al.* 2017). Primarily, sequence-level studies of genome evolution have focussed on single nucleotide polymorphisms and small indels. However, with the advent of long-read sequencing and other technologies that capture long-range linkage information, we are now able to study the effects of larger mutational events such segmental duplications, deletions and other complex structural variants (e.g. Chakraborty *et al.* 2018; Kronenberg *et al.* 2018). Chromosomes may undergo extensive rearrangement via inversions, translocations, fusions and fissions (Eichler and Sankoff 2003). These macro-mutations can have dramatic consequences by altering gene regulation (Farré *et al.* 2019; Stewart and Rogers 2019) and modifying local recombination rates (Farré *et al.* 2013; Martin *et al.* 2019), and they are implicated in key evolutionary processes such as adaptation and speciation (Rieseberg 2001; Kirkpatrick and Barton 2006; Chang *et al.* 2013; Guerrero and Kirkpatrick 2014; Fuller *et al.* 2019; Wellband *et al.* 2019). Chromosome-scale genome assemblies are required to study such macro-mutations, and recent advances in genome assembly have reinvigorated the field (e.g. Dudchenko et al. 2017; Bracewell et al. 2019; Schield et al. 2019; Tandonnet et al. 2019; Bracewell et al. 2020; Teterina et al. 2020). So far, in insects, these studies have been restricted to a few holometabolous genera, such as Diptera (mainly Drosophila and mosquitoes) and Lepidoptera (butterflies) that have been the focus of concerted genome sequencing efforts.

Comparative genomics of Diptera (flies and mosquitoes) and Lepidoptera (butterflies and moths) has revealed high levels of chromosome conservation. For example, tephritid fruit flies have maintained chromosome arms, known as Muller elements (Schaeffer 2018), over at least 60 million years (Sved *et al.* 2016). Conservation of gross chromosome structure is even more striking in mosquitos, where chromosome arms have been maintained for at least 150 million years despite substantial changes in genome size (Dudchenko *et al.* 2017). Among Lepidoptera, the ancestral chromosome complement has largely been maintained over 140 million years, and where changes in karyotype have occurred, they have been driven by chromosome fusion and fission events that maintain ancestral chromosome fragments (d’Alençon *et al.* 2010; Dasmahapatra *et al.* 2012; Ahola *et al.* 2014; Davey *et al.* 2015). The green-veined white butterfly (*Pieris napi*) appears to be one of the few lepidopteran exceptions, as a chromosome-scale reference genome for this insect has recently revealed extensive genome rearrangement despite having a chromosome number similar to model species (Hill *et al.* 2019).

Nonetheless, chromosome number is highly variable across insects as a whole (Blackmon *et al.* 2017), suggesting that the conserved genome structures of Diptera and Lepidoptera cannot be used as models for all insects. A dramatic example of this can be found in aphids – an important group of hemimetabolous sap-sucking plant pests belonging to the insect order Hemiptera – where karyotype varies from 2n = 4 (2 pairs of diploid chromosomes) to 2n = 72 (Blackman 1980). This variation occurs between closely related species, and even within species, suggesting a high rate of chromosome evolution (Blackman 1971; Panigrahi and Patnaik 1991; Blackman *et al.* 2000; Monti *et al.* 2012; Mandrioli *et al.* 2014; Manicardi *et al.* 2015). Aphid chromosome structure and life-cycle may contribute to the rapid evolution of diverse karyotypes (Blackman 1980). Firstly, aphids and other Hemiptera have holocentric chromosomes that lack localised centromeres (Hughes-Schrader and Schrader 1961; Drinnenberg *et al.* 2014). Instead, spindle fibres attach diffusely across the chromosome during meiosis and mitosis (Ris 1942, 1943). As such, both products of a chromosomal fission event can undergo replication, whereas in species with localised centromeres, the fragment lacking the centromere would be lost (Ris 1942; Schrader 1947). Secondly, aphids have an unusual reproductive mode – cyclical parthenogenesis – where they reproduce clonally via apomictic parthenogenesis during the spring, summer and autumn, followed by a sexual stage that produces overwintering eggs that subsequently hatch out as asexually reproducing females (Dixon 1977). Clonal lineages can persist for long periods without sexual reproduction and some species have become obligately asexual (Moran 1992; Simon *et al.* 2002). These bouts of prolonged asexuality, combined with males being derived from an asexual lineage, may enable rearranged karyotypes to persist and potentially contribute to speciation events.

Genome sequencing of a small number of aphid species has also revealed dynamic patterns of genome evolution, with extensive gene duplication having occurred throughout aphid diversification (IAGC 2010; Mathers *et al.* 2017; Thorpe *et al.* 2018; Fernández *et al.* 2019; Julca *et al.* 2019; Li *et al.* 2019). However, the majority of aphid genomes assembled to date are highly fragmented and chromosome-scale genome assemblies have not yet been used to investigate the evolution of diverse aphid karyotypes.

To assess how aphid chromosome architecture compares to that of Diptera and Lepidoptera, we studied chromosome rearrangements of three aphid species spanning approximately 30 million years of aphid evolution. To this end, we generated high-quality chromosome-scale genome assemblies of two extensively studied aphid species: the green peach aphid *Myzus persicae*, a model generalist aphid and major crop pest (Mathers *et al.* 2017); and the pea aphid *Acyrthosiphon pisum*, a model for speciation genomics and basic aphid biology (Hawthorne and Via 2001; Brisson and Stern 2006; Peccoud *et al.* 2009; Pecoud and Simon 2010; Nouhaud *et al.* 2018). Then, we compared these aphid assemblies to a previously published chromosome-scale assembly of the corn-leaf aphid *Rhopalosiphum maidis* (Chen *et al.* 2019). We also analyse the recently released chromosome-scale assemblies of two blood-feeding Hemiptera (*Rhodnius prolixus* and *Triatoma rubrofasciata*), which diverged 60 million ago, and reproduce sexually. We show that there is marked variation across Hemiptera, and between the X and autosomes, in the evolutionary rate of chromosome rearrangement.

## Results and Discussion

### *Chromosome-scale assemblies of the* M. persicae *and* A. pisum *genomes*

High quality, chromosome-scale, genome assemblies of *M. persicae* (clone O) and *A. pisum* (clone JIC1) were generated using a combination of Illumina short-read sequencing, Oxford Nanopore long-read sequencing, 10X Genomics linked-reads (for *A. pisum*) and *in vivo* chromatin conformation capture (HiC) (**Figure 1a** and **b)**. These new genome assemblies provide significant increases in contiguity compared to previously published assemblies at both the contig- and scaffold-levels (**Table 1**; **Supplementary Figure 1**). For *M. persicae*, we report the first chromosome-scale genome assembly of this species with 97% of the assembled content contained in six scaffolds corresponding to the haploid chromosome number of this species (Blackman 1980). Compared to the original assembly of *M. persicae* clone O (Mathers *et al.* 2017), contig number is reduced from 23,616 to 915 and contig N50 is increased by 707% (59 Kb vs. 4.17 Mb). For *A. pisum*, 98% of the assembled content was placed into four scaffolds corresponding to the haploid chromosome number of this species (Blackman 1980). Compared to a recently re-scaffolded reference assembly of *A. pisum* dubbed AL4 (Li *et al.* 2019), we place an additional 14% (98% vs 86%) of the *A. pisum* genome into chromosomes, reduce the number of contigs from 68,186 to 2,298 and increase contig N50 by 1,667% (0.03 Mb vs. 0.53 Mb). K-mer analysis of each assembly versus Illumina short-reads shows very low levels of missing content and the absence of erroneously duplicated content due to the inclusion of haplotigs (allelic variation assembled into separate scaffolds) (**Supplementary Figure 2a** and **b**). Additionally, both assemblies are accurate at the gene-level, containing 94% and 98% of conserved Arthropoda BUSCO genes (n=1,066) as complete, single copies, respectively (**Supplementary Figure 3**). Therefore, the new assemblies of *M. persicae* and *A. pisum* are contiguous, accurate and complete.

**Table 1:** Genome assembly and annotation statistics for *A. pisum, M. persicae* and *R. maidis*. Newly generated assemblies for this study are shaded in grey.

| Species | *A. pisum* | *A. pisum* | *A. pisum* | *M. persicae* | *M. persicae* | *R. maidis* |
| --- | --- | --- | --- | --- | --- | --- |
| Assembly | LSR1 v2 | AL4 v1 | JIC1 v1 | O v1.1 | O v2 | BTI-1 v1 |
| Sequencing approach* | S + IL + MP | HIC** | 10X + ONT + HiC | IL + MP | IL + ONT + HiC | IL + PB + HiC |
| Base pairs (Mb) | 541.68 | 541.12 | 525.80 | 354.7 | 395.14 | 326.02 |
| % Ns | 7.71 | 7.65 | 0.08 | 3.26 | 0.10 | 0.01 |
| Number of contigs | 60,596 | 68,186 | 2,298 | 23,616 | 915 | 960 |
| Contig N50 (Mb)*** | 0.03 | 0.03 | 0.53 | 0.06 | 4.17 | 9.05 |
| Number of scaffolds | 23,924 | 21,919 | 558 | 13,407 | 360 | 220 |
| Scaffold N50 (Mb) | 0.52 | 132.54 | 126.6 | 0.16 | 69.48 | 93.3 |
| % of assembly in chromosome length scaffolds | 0 | 85.96 | 98.20 | 0 | 97.06 | 98.37 |
| Protein coding genes | 36,939 |  | 30,784 | 18,433 | 27,663 | 17,629 |
| Transcripts | 36,939 |  | 34,135 | 30,247 | 31,842 | 17,629 |
| Reference | IAGC (2010) | Li et. al. (2019) | This study | Mathers et. al. (2017) | This Study | Chen et. al. (2019) |

\*S = Sanger, IL = Illumina short reads, MP = Illumina mate-pairs, 10X = 10X genomics linked reads, HiC = high throughput chromatin conformation capture, ONT = Oxford Nanopore long reads, PB = PacBio long reads.
\*\*in vitro (Dovetail Chicago) and in vivo HIC used to correct and scaffold LSR1.
\*\*\*Scaffolds split on runs of 10 or more Ns.

**Figure 1:**
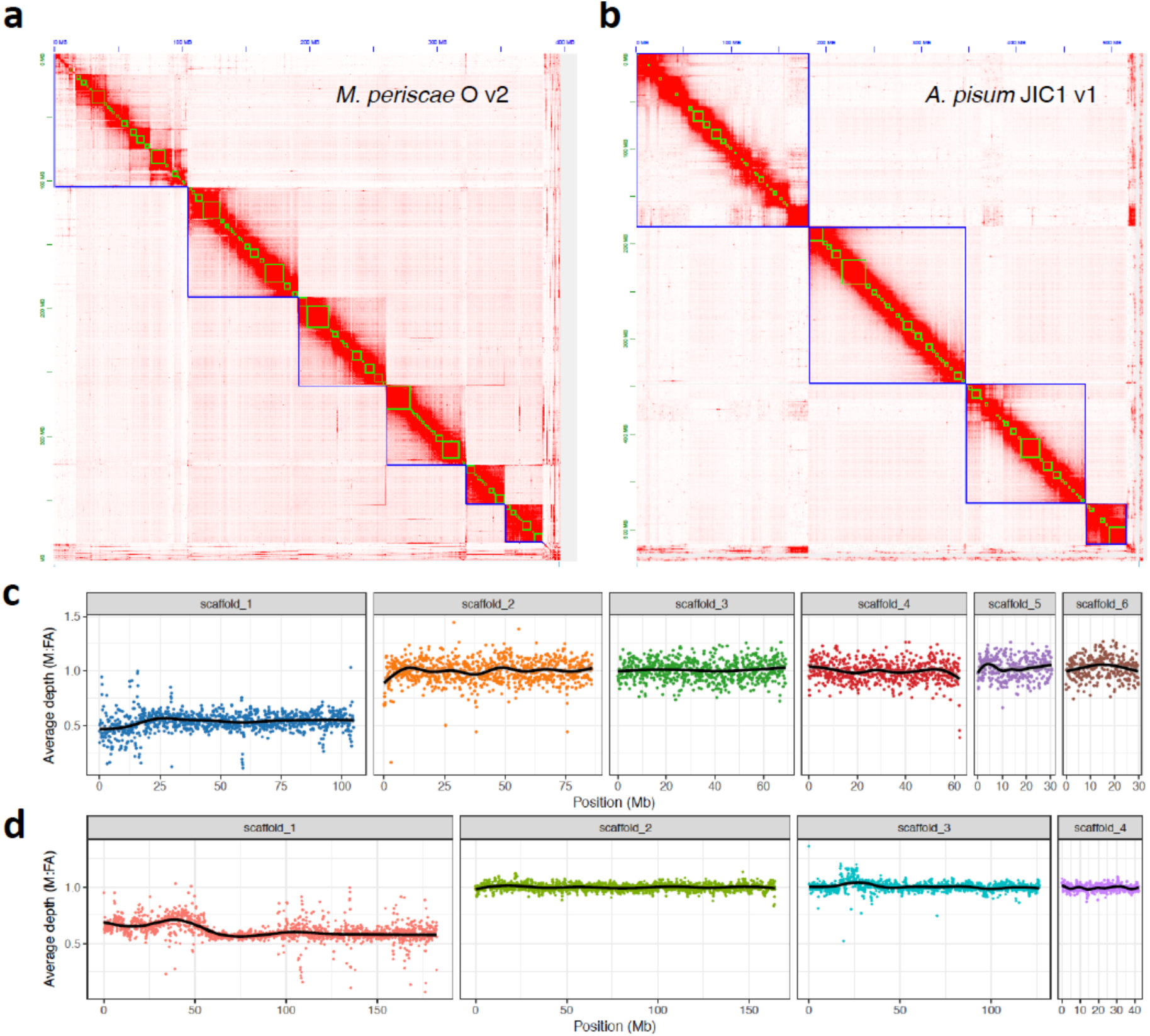
Chromosome-scale genome assemblies of *M. persicae* and *A. pisum*. (**a**) Heatmap showing frequency of HiC contacts along the *M. persicae* clone O v2 (MperO_v2) genome assembly. Blue lines indicate super scaffolds and green lines show contigs. Genome scaffolds are ordered from longest to shortest with the X and Y axis showing cumulative length in millions of base pairs (Mb). (**b**) As for (**a**) but showing HiC contacts along the *A. pisum* JIC1 v1 (ApisJIC1) genome assembly. In this instance, green lines indicate corrected scaffolds from the input assembly which was scaffolded with 10X genomic linked reads prior to chromosome-scale scaffolding with HIC. (**c**) Male (M) to asexual female (FA) coverage ratio of *M. persicae* clone bisulphite sequencing genomic reads in 100kb fixed windows across MperO_v2 chromosome-length scaffolds. The black line indicates the LOESS smoothed average. (**d**) As for (**c**) but showing the male (M) to asexual female (FA) coverage ratio of *A. pisum* clone AL4 genomic reads across ApisJIC1 chromosome-length scaffolds.

Using our improved *M. persicae* and *A. pisum* genome assemblies, we annotated protein coding genes in each species using evidence from RNA-seq data. For *M. persicae*, we aligned 160 Gb of RNA-seq data derived from whole bodies of un-winged asexual females, winged asexual females, winged males and nymphs and annotated 27,663 protein coding genes. For *A. pisum*, we annotated 30,784 protein coding genes, incorporating evidence from 23 Gb of RNA-seq data that were also derived from multiple morphs including un-winged asexual females, sexual females and males. The completeness of the annotations reflected that of the genome assemblies, with 93% and 92% of conserved Arthropoda BUSCO genes (n=1,066) found as complete, single copies, in the *M. persicae* and *A. pisum* annotations, respectively (**Supplementary Figure 4**).

### Identification of the aphid sex (X) chromosome

To identify the X chromosome, we aligned the genomic DNA Illumina reads derived from asexual female and male morphs and calculated the male to asexual female coverage ratio in 100kb fixed windows along each chromosome. Because sex is determined by random loss of one copy of the X chromosome in aphids (Wilson *et al.* 1997), with males carrying a single copy of the X chromosome, males should have half the coverage of females for the X chromosome and equivalent coverage for autosomes (Jaquiéry *et al.* 2018). In agreement with cytological analysis of *M. persicae* and *A. pisum* (Manicardi *et al.* 2014), we find that the longest scaffold in their respective assemblies has the expected coverage pattern of an X chromosome along its full length (**Figure 1c** and **d**). The remaining chromosomes do not deviate from the expected male to asexual female coverage ratio of 1:1, indicating an absence of X chromosome-autosome chimeras. Alignment of *A. pisum* JIC1 with the AL4 assembly and a previously published microsatellite linkage map (Jaquiéry *et al.* 2014) also confirms the identity of the *A. pisum* X chromosome as scaffold 1 and, overall, JIC1 is in broad agreement with AL4 (**Supplementary Figure 5**). Importantly, we assemble and place an additional 50 Mb of the X chromosome in the JIC1 genome assembly compared to AL4, where the X chromosome is only the third longest scaffold and many additional genomic scaffolds with X-chromosome-like coverage patterns are unplaced (Li *et al.* 2019). This is likely due to improved resolution and representation of repetitive elements in JIC1 due to the use of long-read sequence data for de novo assembly. Indeed, for both *M. persicae* and *A. pisum*, we annotate a greater total length of repetitive DNA in our new assemblies than the previous versions that were based on short-read sequencing (*M. persicae* clone O: v1.1 = 57 Mb (16% of total assembly content), v2 = 88 Mb (22%); *A. pisum*: Al4 = 154 Mb (29%), JIC1 = 178 Mb (34%); **Supplementary Figure 6**).

### Extensive autosomal genome rearrangement in aphids

To investigate aphid chromosome evolution, we identified syntenic genomic regions between *M. persicae, A. pisum* and the published chromosome-scale assembly of *R. maidis* (Chen *et al.* 2019) (**Supplementary Table 1**). *M. persicae* and *A. pisum* both belong to the aphid tribe Macrosiphini and diverged approximately 22 million years ago, whereas *R. maidis* belongs to Aphidini and diverged from *M. persicae* and *A. pisum* approximately 33 million years ago (**Figure 2a**). Assessment of chromosomal rearrangements shows a lack of large-scale rearrangements between the X chromosome and the autosomes for any of the aphid species analysed, whereas aphid autosomes have undergone extensive structural change with many rearrangements between chromosomes (**Figure 2c and d**). Comparison between *M. persicae* and *A. pisum* within the tribe Macrosiphini reveals the signature of several chromosome fusion or fission events between autosomes that have occurred within the last 22 million years (**Figure 2c**). For example, *M. persicae* scaffolds 4 and 5 are homologous to *A. pisum* scaffold 3, with the breakpoint clearly delineated. Comparing the more divergent species pair of *M. persicae* and *R. maidis*, which belong to Macrosiphini and Aphidini respectively, reveals highly rearranged autosomes with no clear homology (**Figure 2d**). This is also the case when comparing *R. maidis* to *A. pisum*, despite both species having the same 2n = 8 karyotype (**Supplementary Figure 7**), further supporting high levels of rearrangement. Similar results were obtained by mapping orthologs independently identified based on phylogenomic analysis of gene trees to *M. persicae* chromosomes (**Supplementary Figure 8; Supplementary Table 2a** and **b**). In total we identified 11,372 chromosomally placed one-to-one orthologs between *M. persicae* and *A. pisum* (41% of *M. persicae* genes) and 9,594 between *M. persicae* and *R. maidis* (35% of *M. persicae* genes). Using these data, we confirm that the aphid X chromosome is recalcitrant to translocations, with 93% (1,972 / 2,125) and 96% (1,388 / 1,452) of orthologs conserved on the X chromosome between *M. persicae* and *A. pisum* and between *M. persicae* and *R. maidis*, respectively. Taken together, our results show that the aphid X chromosome has been maintained for at least 33 million years in contrast to extensive autosomal rearrangements.

**Figure 2.**
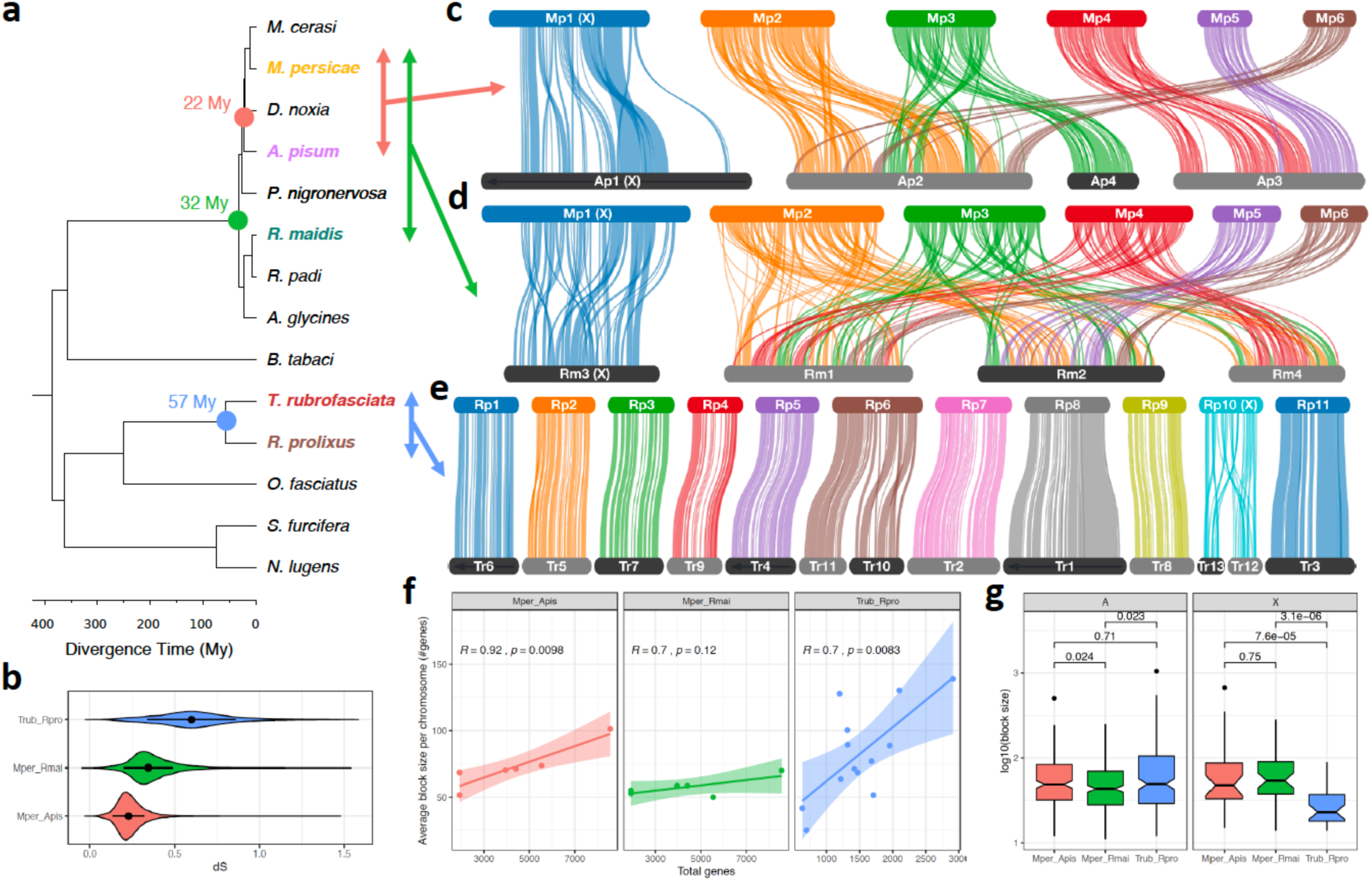
Divergent patterns of chromosome evolution across Hemiptera. (**a**) Time calibrated phylogeny of Hemiptera based on a concatenated alignment of 785 proteins conserved in all species. Divergence times were estimated using non-parametric rate smoothing with calibration nodes specified based on Johnson et. al. (2018). Species with chromosome-scale genome assemblies are coloured and divergence times between focal species are highlighted with coloured circles. (**b**) Synonymous site divergence rate (dS) between *T. rubrofasciata* and *R. prolixus* (blue), *M. persicae* and *R. maidis* (green) and *M. persicae* and *A. pisum* (red) based on 9,087, 7,965 and 9,290 syntenic one-to-one orthologs, respectively. Black circles and whiskers show median and interquartile range respectively. (**c – e**) Pairwise synteny relationships within aphids (**c** and **d)** and Reduviidae (**e**) are mapped onto the phylogeny of Hemiptera. Links indicate the boundaries of syntenic blocks identified by MCscanX and are colour coded by *M. persicae* (**c** and **d**) or *R. prolixus* (**e**) chromosome ID. *A. pisum* (**c**) and *R. maidis* (**d**) chromosomes are ordered based on *M. peircae*, and *T. rubrofasciata* (**e**) chromosomes are ordered according to *R. prolixus*. Black arrows along chromosomes indicate that reverse compliment orientation relative to the focal species. (**f**) The relationship between average synteny block size per chromosome (Y axis) and chromosome size (X axis; measured as the total number of genes per chromosome). Trend lines show linear regression with 95% confidence intervals. For each comparison the Pearson correlation coefficient (R) is given. (**g**) The size of MCscanX synteny blocks (measured in the number of genes within each block) located either on autosomes (A) or the X chromosome (X) for comparisons shown in **c** – **e**. Numbers above comparisons show p values from *Wilcoxon rank-sum tests*.

### Divergent patterns of chromosome evolution across Hemiptera

To investigate how aphid chromosome rearrangements compare to those of other hemipterans, we took advantage of two recently released chromosome-scale assemblies of the blood-feeding species *Rhodnius prolixus* (obtained from the DNA Zoo; Dudchenko et al. 2017) and *Triatoma rubrofasciata* (Liu *et al.* 2019). Both species belong to the hemipteran family Reduviidae and diverged from the aphid lineage approximately 386 million years ago (**Figure 3a**), representing a basal split in extant Hemiptera (Johnson *et al.* 2018). Unlike aphids, most Reduviidae have an XY chromosomal sex determination system (male = XY, female = XX) which is thought to be the ancestral state of Hemiptera (Blackmon *et al.* 2017) and reproduce exclusively through sexual reproduction. In some species, complex sex determination systems have been described with multiple X chromosomes (Ueshima 1966; Panzera *et al.* 1996). *T. rubrofasciata* is one such species and has an X_1_X_2_Y male karyotype (Manna 1950). Multiple X chromosome systems in *Triatoma* are thought to be the result X chromosome fragmentation events (Ueshima 1966), we also examine this hypothesis here.

**Figure 3:**
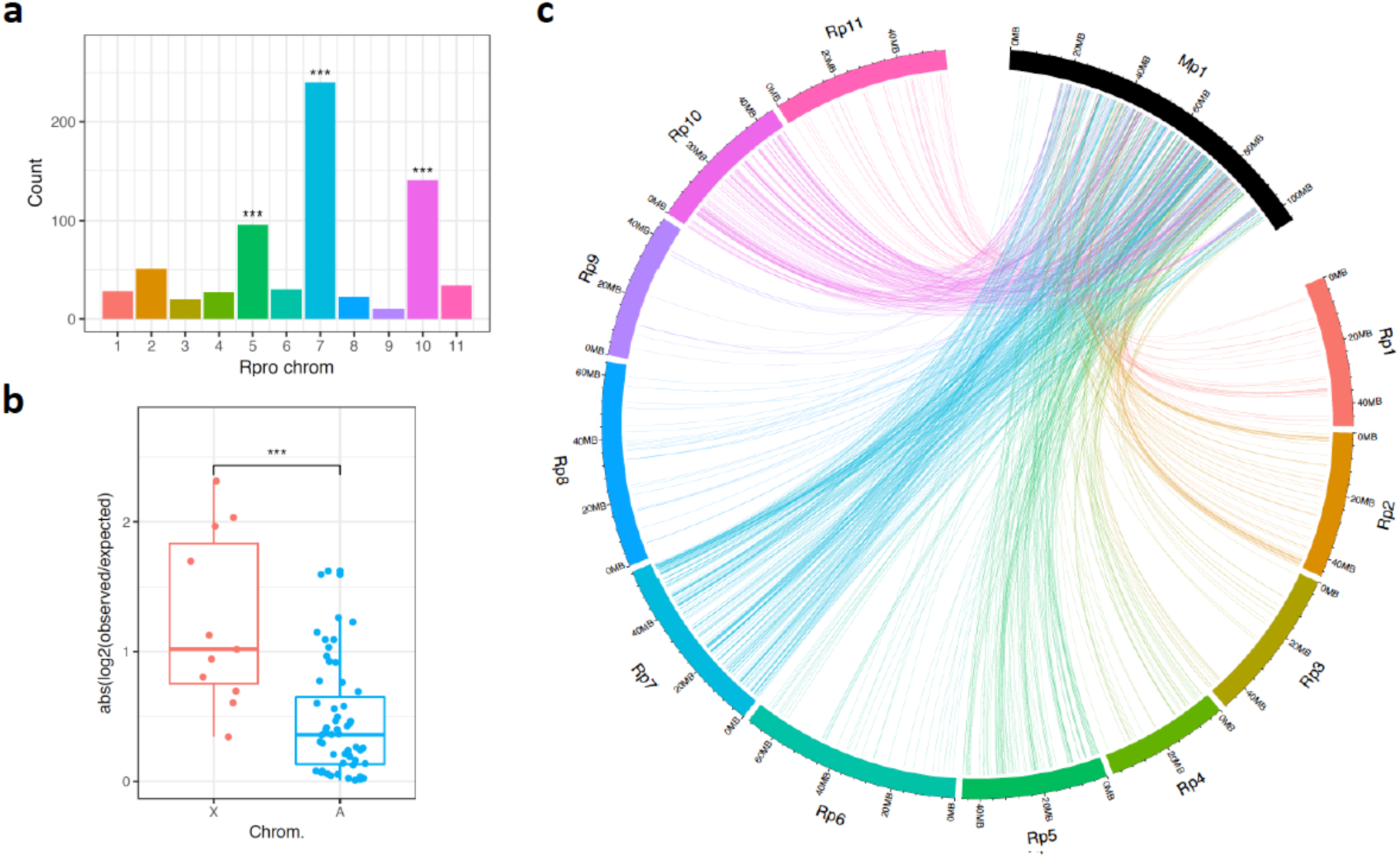
Ortholog mapping between the aphid *Myzus persicae* and the kissing bug *Rhodnius prolixus*. (**a**) Counts of *R. prolixus* chromosomal location for 698 *M. persicae* - *R. prolixus* 1:1 orthologs located on the *M. persicae* X chromosome (scaffold_1). Stars above bars indicate significant enrichment of a specific *R. prolixus* chromosome after correcting for multiple testing (*binomial test*: BH corrected p < 0.05). (**b**) Absolute odds ratios (log_2_(observed/expected) for *R. prolixus* chromosomal enrichment on the *M. persicae* X chromosome and *M. persicae* autosomes. Each dot shows the odds ratio for a specific *R. prolixus* chromosome. *** = *Wilcoxon rank sum test* W = 517, p = 2.31×10^−4^. (**c**) Chord diagram showing links between the *M. persicae* X chromosome (shown as Mp1) and the *R. prolixus* chromosomes for 1:1 orthologs. Rp10 is the *R. prolixus* X chromosome, the *R. prolixus* Y chromosome is not assembled.

In striking contrast to aphids (**Figure 2c** and **d**), *R. prolixus* and *T. rubrofasciata* have highly conserved synteny and an absence of translocation events between chromosomes (**Figure 2e**), despite being almost twice as divergent at the sequence level as the most divergent aphid comparison (**Figure 2b**; median synonymous site divergence: *M. persicae* vs *R. maidis* = 33%, *T. rubrofasciata* vs *R. prolixus* = 60%). In total, just two chromosome fusion or fission events are detectable, one involving *R. prolixus* chromosome 6 (Rp6) and a second involving the X chromosome (Rp10). The latter is likely an X chromosome fission in the *T. rubrofasciata* lineage which has led to the multiple X chromosome sex determination system observed in this species, supporting the hypothesis proposed by Ueshima over half a century ago (Ueshima 1966). For both the *M. persicae* – *A. pisum* comparison and the *T. rubrofasciata* – *R. prolixus* comparison, synteny block size is positively correlated with chromosome length (**Figure 2c**). This relationship breaks down for the *M. persicae* – *R. maidis* comparison, again highlighting high rates of genome rearrangement in aphids. Indeed, despite higher sequence-level divergence, autosomal synteny blocks in Reduviidae are significantly larger than those identified between the most divergent aphid pair of *M. persicae* and *R. maidis* (*Wilcoxon rank-sum test*, W = 19,894, p = 0.02; **Figure 2g**), and are the same size as those identified between the more closely related pair of *M. persicae* and *A. pisum* (*Wilcoxon rank-sum test*, W = 19,086, p = 0.71). This relationship is reversed for synteny blocks on the X chromosome which are significantly larger in aphids than Reduviidae (**Figure 2g**), whether comparing to *M. persicae* – *A. pisum* synteny blocks (*Wilcoxon rank-sum test*: W = 783, p = 7.55 × 10^−5^) or *M. persicae* – *R. maidis* synteny blocks (*Wilcoxon rank-sum test*: W = 1155, p = 3.08 × 10^−6^). Taken together, these results show divergent patterns of both inter- and intra-chromosomal rearrangement rates between aphids and Reduviidae, with dynamic changes in autosome structure potentially having contributed to aphid diversification.

### The hemipteran X chromosome is conserved

To test the hypothesis that the X chromosome is conserved across Hemiptera (Pal and Vicoso 2015) we compared our chromosome-scale assembly of *M. persicae* with *R. prolixus*. We failed to identify syntenic blocks between the two genome assemblies, probably due to the large evolutionary distance between *M. persicae* and *R. prolixus* (386 My). Nonetheless, 6,191 one-to-one orthologs were identified between the two species (22% of *M. persicae* genes), 5,992 (97%) of which are anchored to chromosomes in both species. Using these orthologs, we find that the *M. persicae* X chromosome is significantly enriched for genes located on the *R. prolixus* X chromosome (Rp10) (*binomial test*: BH corrected p = 3.91×10^−13^; **Figure 3a** and **c; Supplementary Figure 9**), suggesting that the aphid and *Rhodnius* X chromosomes are homologous. Furthermore, absolute enrichment (and hence depletion) ratios of orthologs from specific *R. prolixus* chromosomes were significantly higher for the *M. persicae* X chromosome than the autosomes (*Wilcoxon rank sum test*: W = 517, p = 2.31×10^−4^; **Figure 3b; Supplementary Table 3)**, indicating that elevated conservation of the X chromosome, relative to autosomes, extends across Hemiptera. We also find that the *M. persicae* X chromosome is significantly enriched for genes that map to *R. prolixus* autosomes Rp7 (Binomial Test BH corrected p < 1.00×10^−16^) and Rp5 (*binomial test*: Benjamini-Hochberg (BH) corrected p = 3.91×10^−13^) (**Figure 3a** and **c)**. This suggests that he ancestral hemipteran X chromosome may have been fragmented in the *R. prolixus* lineage or, alternatively, the aphid X chromosome may be a product of an ancient chromosome fusion event.

### The aphid X chromosome is repetitive, depleted in expressed genes and rapidly evolving

Conservation of aphid X chromosome gene content is remarkable given its dynamic genomic substrate. In *M. persicae* and *A. pisum*, the X chromosome is significantly more repetitive than the autosomes and significantly depleted in expressed genes (**Figure 4a - d**). Across the *M. persicae* X chromosome, 27% of bases are annotated as TEs compared to 19% in autosomes (*χ*^2^ = 3,486,014, *df* = 1, *p* < 2.2 × 10^−16^). The *A. pisum* X chromosome is even more repetitive, with 42% of bases annotated as TEs compared to 29% in autosomes (*χ*^2^ = 8,455,518, *df* = 1, *p* < 2.2 × 10^−16^). The ends of the X chromosome in both *M. persicae* and *A. pisum* appear to be gene expression deserts with low numbers of expressed genes relative to the autosomes and to the central regions of the X chromosome (**Figure 4a** and **b**). These gene-poor regions have significant reduction in the density of expressed genes towards the telomeres (*M. persicae*: *Pearson correlation* (*R*) = -0.46, p = 6.4 × 10^−7^; *A. pisum*: *R* = -0.46, p = 5.1 × 10^−11^; **Supplementary Figures 10 and 11**). This reduction is associated with significant increases in the densities of DNA transposons (*M. persicae*: *R* = 0.51, p = 1.9 × 10^−8^; *A. pisum*: *R* = 0.63, p < 2.2 × 10^−16^), long terminal repeat (LTR) retrotransposons (*M. persicae*: *R* = 0.52, p = 1.0 × 10^−8^; *A. pisum*: *R* = 0.46, p = 4.4 × 10^−11^), and rolling-circle Helitron transposons (*M. persicae*: *R* = 0.50, p = 6.5 × 10^−8^; *A. pisum*: *R* = 0.38, p = 1.2 × 10^−7^) (**Figure 4a** and **b**; **Supplementary Figures 10 and 11**). There is also a weak but significant increase in long interspersed nuclear elements (LINE) towards the ends of the X chromosome in both species (*M. persicae*: *R* = 0.20, p = 0.04; *A. pisum*: *R* = 0.16, p = 0.029). The age distribution of TE insertions across the genome shows that the invasion of the X chromosome appears to be ongoing, with many young TEs annotated in both *M. persicae* and *A. pisum* (**Figure 4e** and **f**). As well as being repetitive, we also find that genes on the *M. persicae* X chromosome have a higher rate of evolution than those on the autosomes (**Supplementary Figure 12; Supplementary Table 1**), a phenomenon previously observed in *A. pisum* (Jaquiéry *et al.* 2012, 2018). Our results are therefore consistent with a “fast-X” effect operating across aphids. Stability of aphid X chromosome gene content has therefore been maintained in the face of extensive historical, and ongoing, TE activity and high rates of sequence evolution.

**Figure 4:**
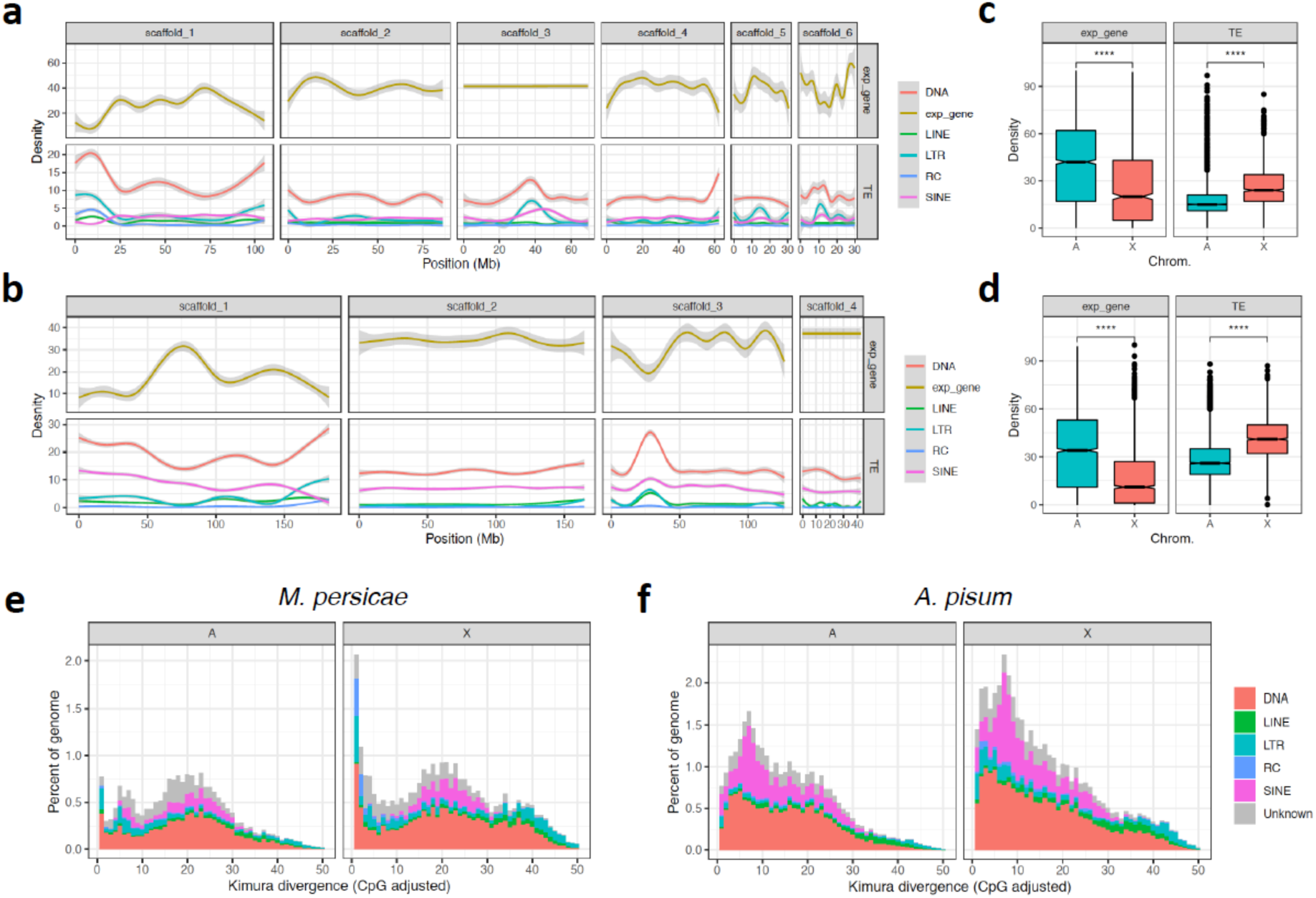
The aphid X chromosome is repetitive and depleted in expressed genes. (**a**) The density of expressed genes (expr_gene) and transposable elements (TEs) across MperO_V2 chromosome-length scaffolds. Genes were classified as expressed if they had an estimated read count > 4 in at least 12 / 24 *M. persicae* morph RNA-seq samples (see **Figure 5**). Lines show LOESS smoothed averages of 100Kb fixed windows. For detailed plots showing all data points for each feature class see **Supplementary Figures 13 and 14**. DNA = DNA transposons, LINE = Long Interspersed Nuclear Elements, LTR = Long Terminal Repeat retrotransposons, RC = Rolling Circle transposons, SINE = Short Interspersed Nuclear Elements. (**b**) As for (**a**) but showing TEs and expressed genes across ApisJIC1 chromosome-length scaffolds. Genes were classified as expressed if they had an estimated read count > 4 in at least 3 / 6 *A. pisum* morph RNA-seq samples from Jaquiéry et al. (2013). (**c**) *Box plots* showing median density of expressed genes and TEs in 100Kb fixed windows across *M. persicae* autosomes and the X chromosome. The X chromosome has significantly lower gene density (*Wilcoxon rank sum test*: W = 1,934,963, p < 2.2×10^−16^) and significantly higher TE density (*Wilcoxon rank sum test*: W = 786,210, p < 2.2×10^−16^) than the autosomes. (**d**) *Box plots* showing median density of expressed genes and TEs in 100Kb fixed windows across *A. pisum* clone JIC1 autosomes and the X chromosome. The X chromosome has significantly lower gene density (*Wilcoxon rank sum test*: W = 16,062,992, p < 2.2×10^−16^) and significantly higher TE density (*Wilcoxon rank sum test*: W = 6,340,780, p < 2.2×10^−16^) than the autosomes. (**e**) *Stacked histograms* showing the age distribution of TEs located on *M. persicae* clone O autosomes (A) and the X chromosome (X). TE families are grouped as for (**a**) and (**b**). (**f**) As for (**e**) but for *A. pisum* clone JIC1.

### Patterns of gene expression along the *M. persicae* genome

Unlike other systems where a fast-X effect is observed (Mank *et al.* 2010), rapid evolution of the aphid X chromosome cannot be explained by reduced efficacy of selection caused by a lower effective population size of the X chromosome relative to autosomes (Jaquiéry *et al.* 2012). This is because progeny produced by aphid sexual reproduction are exclusively female (XX) and inherit an X chromosome from each of their parents, leading to an equivalency of effective population size between the X chromosome and the autosomes (Jaquiéry *et al.* 2012). Rather, aphid fast-X evolution is thought to be predominantly explained by patterns of gene expression. Specifically, lower gene expression levels of X-linked genes compared to those on the autosomes, and enrichment of genes expressed in rare morphs i.e. males and sexual females), possibly driven by antagonistic selection (Jaquiéry *et al.* 2018). Both of these factors lead to relaxed purifying selection on X-linked genes. We examined these hypotheses using our new chromosome-scale assembly of *M. persicae* and a large gene expression data set for diverse *M. persicae* morphs. In particular, we investigated genome-wide patterns of gene expression in un-winged asexual females, winged asexual females, winged males and un-winged asexual female nymphs (**Figure 5a**). We identified 5,046 differentially expressed genes between *M. persicae* morphs assuming a 5% false discovery rate (*Sleuth likelihood ratio test* q < 0.05, absolute effect size (beta) > 0.5 relative to asexual female morphs; **Supplementary Table 4**). Amongst morph-biased genes, 539 are specifically upregulated in males relative to the common wingless asexual female morph (**Figure 5b**). These male-biased genes are significantly enriched on the *M. persicae* X chromosome (*binomial test*: p = 2.38×10^− 6^; **Figure 5c**). Using gene expression data for asexual females, we confirm that the X chromosome has significantly lower gene expression than the autosomes (*Wilcoxon rank-sum test*: W = 715,820, p < 2.2×10-16; **Figure 5d**) and that this is particularly pronounced for the 5’ and 3’ ends of the chromosome (**Figure 5e**).

**Figure 5:**
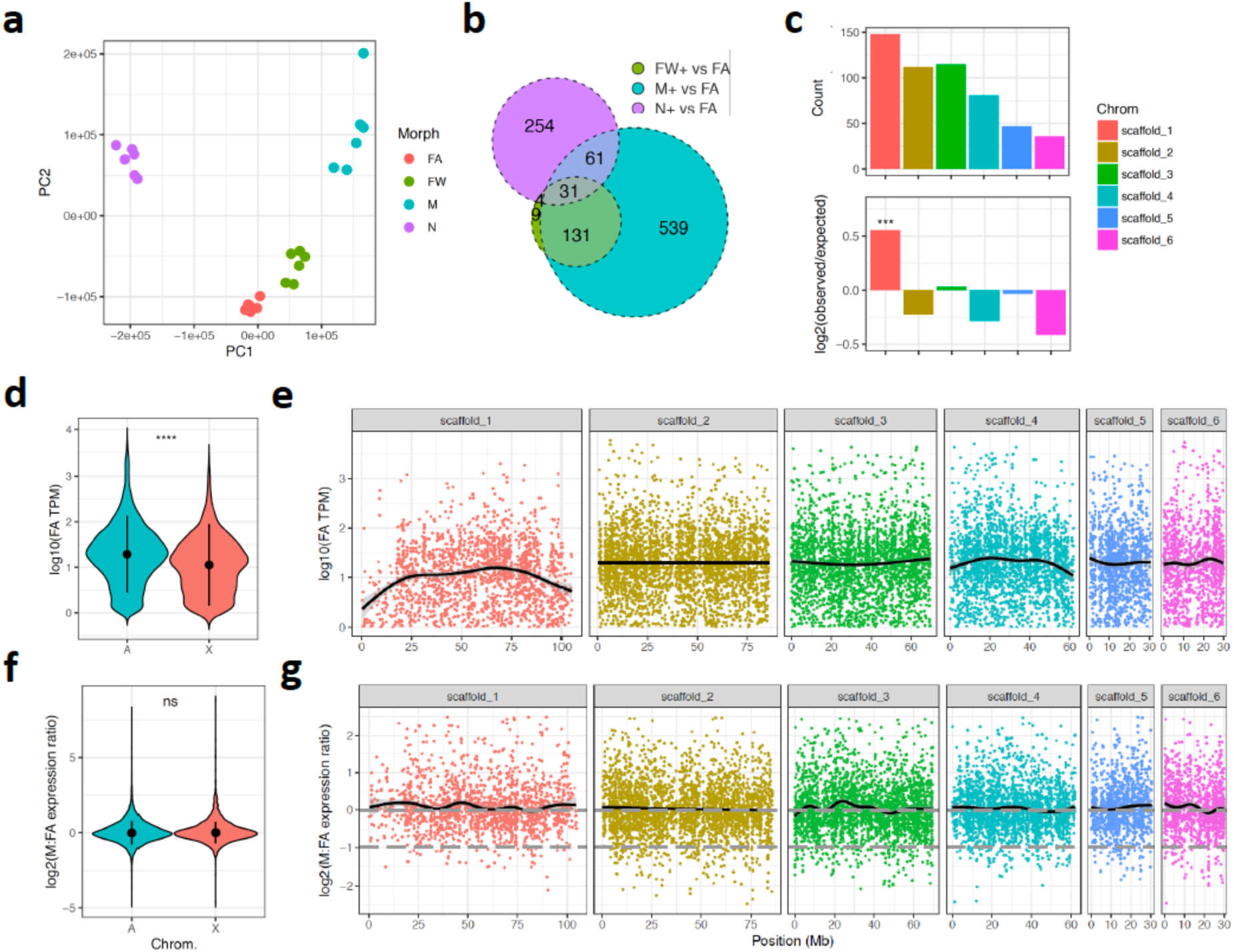
Patterns of gene expression along *M. persicae* chromosomes. (**a**) Principle component analysis (PCA) based on RNA-seq gene expression levels in whole bodies of *M. persicae* clone O un-winged asexual females (FA), winged asexual females (FW), winged males (M) and nymphs (N). Each morph has a distinct gene expression profile with tight clustering of replicates (n=6 per morph). (**b**) Overlap of genes upregulated in either M, FW or N relative to FA (*Sleuth likelihood ratio test*: q < 0.05, effect-size (beta) > 0.5). (**c**) The distribution of genes specifically upregulated in males (n=539) across *M. persicae* clone O v2 chromosome-length scaffolds. Top panel shows counts of M-biased genes per scaffold. Bottom panel shows enrichment scores (log_2_(observed/expected)) of M-biased genes per scaffold relative to the total number of expressed genes on each scaffold (estimated read count > 4 in at least 12 / 24 RNA-seq samples). Significant enrichment was assessed using a binomial test (p < 0.05) with the number of trials equal to the count of expressed genes per scaffold and the probability of success equal to the overall proportion of M-biased genes located on chromosomes relative to the number of expressed genes on all chromosomes. Only the X chromosome is significantly enriched for M-biased genes. *** = *binomial test* p = 2.38×10^−6^. (**d**) *Violin plots* showing the distribution of log_10_ average expression levels (measured in TPM) in FA of expressed genes (TPM > 1) located on *M. persicae* autosomes (A) and the X chromosome (X). The X chromosome has significantly lower gene expression levels than the autosomes (*Wilcoxon rank-sum test*: W = 715,820, p < 2.2×10^−16^). (**e**) FA gene expression ratios used in (**d**) across MperO_v2 chromosome-length scaffolds. Each dot corresponds to a gene, the black line shows the LOESS smoothed average. (**f**) *Violin plots* showing the distribution of log_2_ M to FA gene expression ratios on *M. persicae* autosomes (A) and the X chromosome (X) for genes with average expression of at least 1 TPM in M and FA. Black circles and lines within the coloured regions indicate the median an interquartile range, respectively. There is no significant difference between A and X (*Wilcoxon rank-sum test*: W = 8,919,400, p = 0.10). (**g**) The distribution of log_2_ M to FA gene expression ratios used in (**f**) across MperO_v2 chromosome-length scaffolds. Each dot corresponds to a gene, the black line shows the LOESS smoothed average. The dashed grey lines indicate the expected M to FA gene expression ratio given full dosage compensation (log_2_(M:FA) expression = 0) and in the absence of dosage compensation (log_2_(M:FA) expression = 0.5). Extremely M-biased or FA-biased genes (abs. log_2_ M:FA expression ratio > 2.5) are excluded.

Finally, we also confirm the operation of dosage compensation in *M. persicae*, with no overall difference observed in the male to asexual female gene expression ratio between the X chromosome and the autosomes, despite the X chromosome being found as a single copy in males (*Wilcoxon rank-sum test*: W = 8,919,400, p = 0.10; **Figure 5f**; **Supplementary Table 5**). Dosage compensation has previously been shown to operate in other Hemiptera (Pal and Vicoso 2015) and in *A. pisum* (Jaquiéry *et al.* 2013; Richard *et al.* 2017) using fragmented assemblies. Using our new chromosome-scale assembly of *M. persicae*, we are able to show that dosage compensation operates across the entire X chromosome (**Figure 5f**).

## Conclusion

Aphids show extensive autosomal genome rearrangements. This is in contrast to other insect genomes that have been compared thus far, including within Lepidoptera and Diptera. Furthermore, the high rate of autosomal rearrangements does not appear to be a ubiquitous feature of Hemiptera given that two other Hemiptera (*R. prolixus* and *T. rubrofasciata*) have highly conserved synteny (**Figure 2e**). Our data supports previous karyotype studies showing that chromosome numbers are high variable among aphids (Blackman 1980). Furthermore, our data reveal that the chromosome number variation is not only caused by chromosome fission or fusion (i.e. macro-mutations), but also by autosomal recombination events.

In contrast, the aphid X chromosome appears to recalcitrant to rearrangement with the autosomes, and it appears structurally highly conserved. The long-term stability of aphid X chromosome gene content is surprising, given that we observed low levels of gene expression of X-linked genes, relaxed selection on the coding genes, and an accumulation of transposable elements. This implies that strong selection is acting against inter-chromosomal rearrangements involving the X chromosome. Selection is likely to operate during the sexual phase of the aphid life cycle as X chromosome fissions have been observed in *M. persicae* populations (Monti *et al.* 2012; Mandrioli *et al.* 2014). One possible explanation is that large-scale translocations involving the X chromosome interfere with dosage compensation, causing the mis-expression of genes. Alternatively, intact X chromosomes may be required for proper elimination of the X chromosome during male determination. Altogether, this study shows that long-read sequencing and chromosome-scale assemblies can uncover macro-mutations and non-homologous recombination events that are likely to have significantly impacted evolution. Aphids serve as an excellent model system to understand the role of such genome rearrangements in evolution.

## Methods

### Aphid genome assembly strategy

To assemble high-quality reference genomes for *M. persicae* and *A. pisum*, we generated initial *de novo* contig assemblies based on high-coverage Nanopore long-read data. These assemblies were then scaffolded into pseudomolecules (chromosomes) using *in vivo* chromatin conformation capture (HiC) data (Dudchenko *et al.* 2017) and, in the case of *A. pisum*, 10x Genomics Chromium linked-reads (Zheng *et al.* 2016; Weisenfeld *et al.* 2017). As *M. persicae* and *A. pisum* have divergent genome architectures (e.g. repeat content and level of heterozygosity), we optimised the initial contig assembly for each species, aiming to maximise genome completeness and minimise pseudo duplication caused by under-collapsed heterozygosity. These criteria were assessed by comparing the K-mer content of raw sequencing reads to the genome assembly with the K-mer Analysis Toolkit (KAT) (Mapleson *et al.* 2017) and by assessing the representation of conserved genes with BUSCO v3 (Simão *et al.* 2015; Waterhouse *et al.* 2018), using the Arthropoda gene set (n=1,066). We also used genome size estimates for *M. persicae* (409 Mb) and *A. pisum* (514 Mb) based on flow cytometry from Wenger et. al. (2017) to assess the proportion of the genome that had been assembled and to estimate sequence read coverage. For each species, we compared long-read assemblies generated with Canu (Koren *et al.* 2017), Flye (Kolmogorov *et al.* 2019) and wtdbg2 (Ruan and Li 2019) as well as various combinations of assembly merging with quickmerge (Chakraborty *et al.* 2016), the effect of removing alternative haplotypes and the effect of long- and short-read assembly polishing (**Supplementary Note**). Below, we describe the steps used to generate the final genome assembly for each species.

### *Sequencing and* de novo *assembly of* M. persicae *clone O*

We previously sequenced the genome of *M. persicae* clone O using Illumina short-read sequencing (Mathers *et al.* 2017). We used aphids derived from the same asexually reproducing colony maintained at the John Innes Centre insectary for all DNA extractions.

For Nanopore long-read sequencing, batches of twenty aphids were collected in 1.5 ml low-bind Eppendorf tubes and snap-frozen in liquid nitrogen. We extracted high molecular weight DNA with the Illustra Nucleon PhytoPure kit (GE Healthcare, RPN8511) following the manufacturers protocol. Wide-bore pipette tips were used when transferring solutions to circumvent shearing of DNA. DNA concentration was determined using the Qubit broad-range assay. The purity of each extraction was assessed using a NanoDrop spectrophotometer (Thermo Fisher) based on 260/280 nm and 260/230 nm absorbance values, and by comparing the NanoDrop concentration estimate to the Qubit estimate, looking for a ratio close to 1:1 (Schalamun *et al.* 2019). The length of extracted DNA molecules was assessed using a Femto fragment analyser (Agilent). Nanopore genomic DNA libraries were prepared for samples passing quality control using the Ligation Sequencing Kit (Oxford Nanopore Technologies (ONT), Oxford, UK: SQK-LSK109) following the manufacturers protocol with the exception that we started with 10 µg of high molecular weight DNA. In total, four libraries were generated and each one sequenced on an R9.4 flow cell for 72 hours. Base-calling was run using Guppy v2.3.1 (ONT, Oxford, UK) with default settings, retaining reads with a quality score of at least This resulted in a total of 28 Gb of data (∼70x coverage of the *M. persicae* genome) with a an N50 of 23 Kb (**Supplementary Table 6**).

We also generated 24 Gb (∼59x coverage) of Illumina short-reads for assembly polishing and quality control. DNA was extracted from ∼50 individuals with a modified CTAB protocol (Marzachi *et al.* 1998) and sent to Novogene (China) for sequencing. Novogene prepared a PCR-free Illumina sequencing library using the NEBNext Ultra II DNA Library Prep Kit for Illumina (New England Biolabs, USA), with the manufacturers protocol modified to give a 500 bp – 1 kb insert size. This library was sequenced on an Illumina HiSeq 2500 instrument with 250 bp paired-end chemistry. The resulting reads were trimmed for adapter sequences with trim_galore! v0.4.0 (Krueger 2015), retaining read pairs where both sequences were at least 150 bp long after adapter trimming.

In our exploratory analysis, wtdgb2 v2.3 gave optimum performance for assembling the *M. persicae* clone O Nanopore data (**Supplementary Note**). We generated two wtdgb2 assemblies with the parameters “-x ont -p 0 -k 17 -L 15000” and “-x ont -p 19 -k 0 -L 15000”. These assemblies had complementary contiguity and contained non-overlapping sets of BUSCO genes. We therefore merged the two wtdgb2 genome assemblies with quickmerge v0.3 using the parameters “-l 1837291 -ml 10000”, with the more complete wtdgb2 “-x ont - p 0 -k 17 -L 15000” assembly used as the query. This resulted in an assembly that was more complete and more contiguous than either individual wtdgb2 assembly (see **Supplementary Note)**. The merged wtdgb2 assembly was then iteratively polished, first with three rounds of long-read polishing with racon v1.3.1 (Vaser *et al.* 2017), then with three rounds of short-read polishing with Pilon v1.22 (Walker *et al.* 2014) in diploid mode. Redundant haplotigs (contigs derived from un-collapsed heterozygosity) were removed from the polished assembly with Purge Haplotigs (Roach *et al.* 2018) using the sequence coverage bounds 9, 45 and 92, and requiring contigs to cover at least 90% of another, longer contig, to be flagged as a haplotig.

### *Sequencing and* de novo *assembly of* A. pisum *clone JIC1*

An isolate of *A. pisum* (dubbed JIC1) found on *Lathyrus odoratus* (sweet pea) was collected from Norwich in 2005 and subsequently reared at the JIC insectary under controlled conditions (Dr Ian Bedford, personal communication). DNA extractions and Nanopore sequencing libraries were prepared as described above for *M. persicae* clone O. In total, two libraries were generated and each one sequenced on an R9.4 flow cell for 72 hours. Base calling was run using Guppy v2.3.1 with the “flip-flop” model, retaining reads with a quality score of at least 7. This resulted in a total of 18 Gb of data (∼35x coverage of the *A. pisum* genome) with an N50 of 33 Kb (**Supplementary Table 6**).

To improve the Nanopore *de novo* assembly and generate accurate Illumina short-reads for assembly polishing, we generated 10X Genomics Chromium linked-read data using DNA extracted as described above. High molecular weight DNA was sent to Novogene (China) for 10X Genomics Chromium library preparation following the manufacturers protocol and sequencing was performed on an Illumina NovaSeq instrument. In total we generated 45 Gb of 150 bp paired-end reads (∼88x coverage of the *A. pisum* genome). The average molecule size of the library was 32 Kb (**Supplementary Table 6**).

*De novo* assembly with Flye v2.4 using default settings gave the best balance between contiguity, genome completeness and absence of erroneously duplicated content (**Supplementary Note**). The Flye assembly was polished as described above for *M. persicae*, with three rounds of racon followed by three rounds of Pilon. For Pilon polishing, we used the 10X reads after removing barcodes and primer sequence with process_10xReads.py (https://github.com/ucdavis-bioinformatics/proc10xG). Redundant haplotigs were removed from the polished Flye assembly with Purge Haplotigs (Roach *et al.* 2018) using the sequence coverage bounds 4, 21 and 57, and requiring contigs to cover at least 75% of another, longer contig, to be flagged as a haplotig. Finally, we iteratively scaffolded the de-duplicated Flye assembly using our 10X Genomics linked-read data. We ran two iterations of Scaff10x v4.0 (https://github.com/wtsi-hpag/Scaff10X) with the parameters “-longread 1 -edge 45000 - block 45000” followed by Tigmint v1.1.2 (Jackman *et al.* 2018) with default settings, which identifies misassemblies, breaks the assembly and performs a final round of scaffolding with ARCS (Yeo *et al.* 2018).

### HiC libraries and genome scaffolding

To scaffold our *de novo* assemblies of *M. persicae* clone O and *A. pisum* clone JIC1 we used *in vivo* chromatin conformation capture to generate HiC data. For each species, whole bodies of individuals from the same clonal populations used for genome sequencing were snap frozen in liquid nitrogen and sent to Dovetail Genomics (Santa Cruz, California, USA) for HiC library preparation and sequencing. HiC libraries were prepared using the *DpnII* restriction enzyme following a similar protocol to Lieverman-Aiden et al. (2009). HiC libraries were sequenced on an Illumina HiSeq X instrument, generating 150 bp paired-end reads. In total, we generated 123 Gb (∼300x coverage) and 21 Gb (∼40x coverage) of HiC data for *M. persicae* clone O and *A. pisum* clone JIC1, respectively (**Supplementary Table 6**). To identify HiC contacts, we aligned our HiC data to our draft assemblies using the Juicer pipeline (Durand *et al.* 2016). We then used the 3D-DNA assembly pipeline (Dudchenko *et al.* 2017) to first correct misassemblies in each input assembly and then to order contigs (or scaffolds for *A. pisum* JIC1) into superscaffolds. K-mer analysis showed that our draft assemblies did not contain substantial quantities of duplicated content caused by the inclusion of haplotigs so we ran 3D-DNA in “haploid mode” with default settings for *M. persicae* clone O and “--editor-repeat-coverage 4” for *A. pisum* JIC1 (**Supplementary Note**). The initial HiC assembly for each species was then manually reviewed using Juicebox Assembly Tools (JBAT) to correct misjoins and other errors (Dudchenko *et al.* 2018). Following JBAT review, the assemblies were polished with the 3D-DNA seal module to reintegrate genomic content removed from superscaffolds by false positive manual edits to create a final scaffolded assembly. The HIC assemblies were then screened for contamination with BlobTools (Kumar *et al.* 2013; Laetsch and Blaxter 2017). Finally, a frozen release was generated for each assembly with scaffolds renamed and ordered by size with SeqKit v0.9.1 (Shen *et al.* 2016). The final assemblies were checked with BUSCO and KAT comp to ensure the scaffolding and decontamination steps had not reduced gene-level completeness or removed genuine single-copy aphid genome content.

### *Transcriptome sequencing of* M. persicae *morphs*

We previously sequenced the transcriptomes *M. persicae* clone O apterous (un-winged) asexual females and alate (winged) males using six biological replicates per morph (Mathers *et al.* 2019). As part of the same experiment we also collected and sequenced nymphs (derived from apterous asexual females) and alate asexual females (also six biological replicates each). These data were not used in our original study (Mathers *et al.* 2019) but are included here for genome annotation and to provide a more comprehensive view of morph-biased gene expression in *M. persicae*. Aphid rearing, RNA extraction and sequencing were carried out as in Mathers et. al. (2019). Apterous asexual females, alate asexual females and nymphs were reared in long day conditions (14 hr light, 22°C day time, and 20°C night time, 48% relative humidity) and alate males were reared in short day conditions (8 hr light, 18°C day time, and 16°C night time, 48% relative humidity).

### Genome annotation

We annotated protein coding genes in our new chromosome-level assemblies of *M. persicae* and *A. pisum* using BRAKER2 v2.1.2 (Hoff *et al.* 2015, 2019), incorporating evidence from RNA-seq alignments. Prior to running BRAKER2, we soft-masked each genome with RepeatMasker v4.0.7 (Tarailo-Graovac and Chen 2009; Smit *et al.* 2015) using known Insecta repeats from Repbase (Bao *et al.* 2015) with the parameters “-e ncbi -species insecta -a -xsmall -gff”. We then aligned RNA-seq data to the soft-masked genomes with HISAT2 v2.0.5 (Kim *et al.* 2015). All RNA-seq data sets used for annotation are summarised in **Supplementary Table 7**. For *M. persicae*, we aligned 25 RNA-seq libraries. Specifically, we used a high coverage (∼200 million reads), strand-specific, RNA-seq library generated from mixed whole bodies of apterous *M. persicae* clone O asexual females (Mathers *et al.* 2017) as well as newly generated (see above) and publicly available (Mathers *et al.* 2019) un-stranded RNA-seq data for *M. persicae* clone O nymphs (derived from apterous asexual females), alate asexual females, apterous asexual females and males (six biological replicates each). All RNA-seq data was trimmed for adapters and low quality bases (quality score < 20) with Trim Golore v0.4.5 (Krueger 2015), retaining reads where both members of the pair are at least 20bp long. Un-stranded RNA-seq data was aligned to the genome with HISAT2 with the parameters “--max-intronlen 25000 --dta-cufflinks” followed by sorting and indexing with SAMtools v1.3 (Li *et al.* 2009). Strand-specific RNA-seq was mapped as for the un-stranded data, with the addition of the HISAT2 parameter “--rna-strandness RF”. We then ran BRAKER2 with UTR training and prediction enabled with the parameters “--softmasking --gff3 --UTR=on”. Strand-specific RNA-seq alignments were split by forward and reverse strands and passed to BRAKER2 as separate BAM files to improve the accuracy of UTR models as recommended in the BRAKER2 documentation. For *A. pisum* clone JIC1, we used un-stranded RNA-seq data derived from whole bodies of *A. pisum* clone LSR1 (IAGC 2010) males, asexual females and sexual females (two biological replicates each) from Jaquiéry et al. (2013). Reads were, trimmed, mapped and passed to BRAKER2 as for the un-stranded *M. persicae* RNA-seq data. Following gene prediction, genes were removed that contained in frame stop codons using the BRAKER2 script getAnnoFastaFromJoingenes.py and the completeness of each gene set was checked with BUSCO using the longest transcript of each gene as the representative transcript.

### X chromosome identification

We identified the aphid sex (X) chromosome in our new assemblies of *M. persicae* clone O and *A. pisum* JIC1 based on the ratio of male (M) to asexual female (FA) coverage of Illumina genomic DNA reads. For *M. persicae*, we used whole genome bisulphite sequencing (BS-seq) reads from Mathers et al. (2019), merging biological replicates by morph. These data are derived from the same clonal population (clone O) as used for the genome assembly. BS-seq reads were aligned to the *M. persicae* clone O v2 genome with Bismark v0.20.0 (Krueger and Andrews 2011) with default parameters. We used Sambamba v0.6.8 to estimate BS-seq read depth in 100 Kb fixed windows for M and FA separately using the BAM files generated by Bismark and the parameters “depth window --fix-mate-overlaps --window-size=100000 -- overlap=100000”. We then calculated the ratio of M to FA read depth per window (i.e. the coverage ratio). Coverage ratios showed scaffold 1 to have the expected X chromosome M to FA coverage ratio (50% that of the autosomes). To generate **Figure 1c** we calculated average M (107x) and FA (82x) coverage excluding scaffold 1 to derive a coverage correction factor for FA (x1.3), and used this to calculate normalised M to FA coverage ratio for each 100 Kb window. For *A. pisum* JIC1, we used whole genome Illumina sequence data of clone AL4 M and FA morphs the from Li et al. (2019). We followed the same procedure as for *M. persicae* clone O with the exception of using BWA-MEM v0.7.17 (Li 2013) to map reads and Sambamba markdup to identify reads derived from PCR duplicates prior to calculating coverage statistics. Scaffold 1 was identified as the X chromosome. Excluding scaffold 1, we calculated average M (45x) and FA (41x) coverage to derive a coverage correction factor for FA (x1.1), and used this to calculate normalised M to FA coverage ratio for each 100 Kb window along *A. pisum* JIC1 chromosome length scaffolds to generate **Figure 1d**.

### *Re-annotation of the chromosome-scale assemblies of* R. prolixus *and* T. rubrofasciata

We included the recently released chromosome-scale genome assemblies of the blood-feeding hemipterans *Rhodnius prolixus* (obtained from the DNA Zoo (https://www.dnazoo.org/); Dudchenko et al. 2017) and *Triatoma rubrofasciata* (Liu *et al.* 2019) in our synteny and phylogenomic analyses. The *R. prolixus* chromosome-level assembly has not yet been annotated and we found on initial inspection that the *T. rubrofasciata* gene release is based on the contig assembly of this species and not the chromosome-length scaffolds. We therefore generated *de novo* gene predictions for these two species using BRAKER2 with evidence from protein alignments created with GenomeTheader v1.7.1 (Gremme 2014). For each species, we soft-masked the genome for known repeats as for *M. persicae* and *A. pisum*. We then ran BRAKER2 with the parameters “--softmasking --gff3 -- prg=gth --trainFromGth”. For *R. prolixus*, we used proteins from the original gene release as evidence (Mesquita *et al.* 2015). For *T. rubrofasciata* we used proteins from Liu et al. (2019). The final BRAKER2 gene sets for each species were checked completeness using BUSCO as for *M. persicae* and *A. pisum*.

### Phylogenomic analysis of sequenced hemipteran genomes

We estimated a time calibrated phylogeny of Hemiptera using protein sequences from our new genome assemblies of *M. persicae* clone O and *A. pisum* clone JIC1, the new annotations of the chromosome-scale assemblies of *R. prolixus* and *T. rubrofasciata* and ten previously sequenced Hemiptera: *Myzus cerasi* (Thorpe *et al.* 2018), *Diuraphis noxia* (Nicholson *et al.* 2015), *Pentalonia nigronervosa* (Mathers *et al.* in prep.), *Rhopalosiphum maidis* (Chen *et al.* 2019), *Rhopalosiphum padi* (Thorpe *et al.* 2018), *Aphis glycines* (version 2) (Mathers 2020), *Bemisia tabaci* MEAM1 (Chen *et al.* 2016), *Oncopeltus fasciatus* (Panfilio *et al.* 2019), *Sogatella furcifera* (Wang *et al.* 2017) and *Nilaparvata lugens* (Xue *et al.* 2014). Where multiple transcripts of a gene were annotated we used the longest transcript to represent the gene model. We used OrthoFinder v2.2.3 (Emms and Kelly 2015, 2019) with Diamond v0.9.14 (Buchfink *et al.* 2014), MAFFT v7.305 (Katoh and Standley 2013) and FastTree v2.1.7 (Price *et al.* 2009, 2010) to cluster proteins into orthogroups, reconstruct gene trees and estimate the species tree. The OrthoFinder species tree was rooted according to Johnson et al. (2018). To estimate approximate divergence times for our taxa of interest, we used penalised likelihood implemented in r8s with secondary calibration points derived from Johnson et al. (2018) (**Supplementary Table 8**).

### Synteny analysis

We identified syntenic blocks of genes between *M. persicae, A. pisum* and *R. maidis*, and between *R. prolixus* and *T. rubrofasciata*, using MCScanX v1.1 (Wang *et al.* 2012). For each comparison, we carried out an all vs. all BLAST search of annotated protein sequences using BLASTALL v2.2.22 (Altschul *et al.* 1990) with the options “-p blastp - e 1e-10 -b 5 -v 5 -m 8” and ran MCScanX with the parameters “-s10 -b 2”, requiring synteny blocks to contain at least ten consecutive genes and to have a gap of no more than 25 genes. MCScanX results were visualised with SynVisio (https://synvisio.github.io/#/). We parsed the MCScanX results and estimated synonymous and nonsynonymous substitution rates between pairs of syntenic genes using collinearity scripts from Nowell et al. (2018; https://github.com/reubwn/collinearity). We also investigated synteny using orthologous genes identified by OrthoFinder. We performed two additional OrthoFinder runs, one with the chromosome-scale assemblies of *M. persicae, A. pisum* and *R. maidis*, and one using the three aphid assemblies and the chromosome-scale assembly of *R. prolixus*. OrthoFinder was run as described above for the phylogenomic analysis of Hemiptera.

To test for conservation of the X chromosome across Hemiptera, we first identified *R. prolixus* chromosomes that were likely to be homologous to *M. persicae* chromosomes. We therefore mapped their orthologous genes onto chromosomes. Next, we tested for significant enrichment of genes from specific *R. prolixus* (target) chromosomes on each *M. persicae* (focal) chromosome using a *binomial test.* In each *binomial test*, the observed ortholog count from a target *R. prolixus* chromosome is the *number of successful trials.* The total number of orthologs on the *M. persicae* focal chromosome is the *total number of trials* (this is equal to the sum of all *R. prolixus* orthologs that map to the focal chromosome). Finally, the *probability of success* is equal to the faction orthologs found on the *R. prolixus* target chromosome, relative to the total number of orthologs. We corrected for multiple testing using the Benjamini-Hochberg (BH) procedure (Benjamini and Hochberg 1995). For each focal *M. persicae* chromosome we also calculated the observed / expected ratio of orthologs from each target *R. prolixus* chromosome. The expected ortholog count was calculated by multiplying the total ortholog count for the focal *M. persicae* chromosome by the faction of all *M. persicae* – *R. prolixus* orthologs found on the target *R. prolixus* chromosome.

### M. persicae *gene expression*

We investigated patterns of gene expression in the *M. persicae* clone O v2 genome using newly generated (see above) and previously published (Mathers *et al.* 2019) RNA-seq data for *M. persicae* clone O nymphs (derived from un-winged asexual females), winged asexual females, un-winged asexual females and winged males (six biological replicates each). Transcript-level expression was estimated for each sample with Kallisto v0.44.0 (Bray *et al.* 2016) with 100 bootstrap replicates. We identified differentially expressed genes between *M. persicae* morphs using Sleuth (Pimentel *et al.* 2017), aggerating transcript-level p values (Yi *et al.* 2018). Specifically, we used a likelihood ratio test (LRT) to identify genes that significantly vary by morph (BH corrected p < 0.05). To quantify the magnitude of the change in expression relative to un-winged asexual females (from which the other morphs are derived), we applied pairwise Wald Tests between un-winged asexual females and each alternative morph and recorded the effect size (beta) which approximates the log_2_ fold change in expression. We considered genes to be “morph-biased” if they had a significant LRT result and abs. beta > 0.5 in any morph relative to un-winged asexual females. To identify genes that were specifically up-regulated in males, we identified the subset of “morph biased” genes that had beta > 0.5 in winged males and beta < 0.5 in winged asexual females and nymphs.

To test for dosage compensation in *M. persicae* clone O, we calculated the log2 ratio of winged male to un-winged asexual female gene expression using transcripts per million (TPM) expression values estimated by Kallisto for all genes with expression of at least one TPM in both morphs. For each gene, we used the longest transcript to represent the gene. We then compared expression ratios for genes on the X chromosome and the autosomes with a Wilcoxon rank-sum test.

### Transposable element analysis

To investigate the distribution of transposable elements (TEs) in *M. persicae* clone O v2 and *A. pisum* JIC1 v1 we generated a comprehensive TE annotation. For each assembly, we modelled TEs *de novo* with RepeatModeler v1.0.8 (Smit and Hubley 2008) and then merged the *de novo* repeats with known repeats from the RepBase Insecta library (Bao *et al.* 2015) using ReannTE_MergeFasta.pl (https://github.com/4ureliek/ReannTE). We then annotated TEs across each genome with RepeatMasker v4.0.7 (Smit *et al.* 2005; Tarailo-Graovac and Chen 2009) using the species-specific merged TE library. We calculated TE density in 100 Kb and 1 Mb fixed windows with DensityMap (Guizard *et al.* 2016), grouping all TEs together, and also separately for DNA transposons, long interspersed nuclear elements, long terminal repeat retrotransposons, rolling circle transposons and short interspersed nuclear elements. We also calculated the density of expressed genes in the same windows. For *M. persicae*, we used genes classified as expressed by sleuth (estimated count > 4 in at least 12 / 24 samples) in the “morph biased” expression analysis (above). To generate equivalent data for *A. pisum*, we ran Kallisto and Sleuth as for the *M. persicae* morph-biased expression analysis (above) using RNA-seq data derived from whole bodies of *A. pisum* clone LSR1 (IAGC 2010) males, asexual females and sexual females (two biological replicates each) from Jaquiéry et al. (2013). Genes were considered expressed if they had an estimated read count > four in at least three out of six samples.

To generate TE age distributions for *M. persicae* clone O v2 and *A. pisum* JIC1 we ran RepeatMasker separately for the autosomes and the X chromosome for each species and parsed the output with parseRM_GetLandscape.pl (https://github.com/4ureliek/Parsing-RepeatMasker-Outputs). We used the CpG adjusted Kimura 2-parameter distance of each TE insertion from its corresponding consensus sequence as a proxy for TE age.

## Data availability

Raw sequence data generated for this study are available at the NCBI short-read archive under BioProject PRJNA613055. Genome assemblies, annotations and supplementary data are available from Zenodo: https://10.5281/zenodo.3712089.

## Acknowledgements

This work was funded by a BBSRC Future Leader Fellowship (BB/R01227X/1) awarded to TCM, the BBSRC Industrial Partnership Award (IPA) with Syngenta Ltd (BB/L002108/1 and BB/R009481/1) awarded to SAH, DS and CvO and BBSRC PhD fellowship of RW. Additional support was received from the BBSRC Institute Strategy Programme (BB/P012574/1) and the John Innes Foundation. This research was supported in part by the NBI Computing Infrastructure for Science Group, which provides technical support and maintenance to the John Innes Centre’s high-performance computing cluster and storage systems. We thank the JIC Entomology Facility for assistance with rearing of aphids and in particular Dr. Ian Bedford for collecting the isolate of *A. pisum* dubbed JIC1 and Anna Jordan for identification of *M. persicae* morphs.

