## Supplementary Figure for "Chromosome-scale genome assemblies of aphids reveal extensively rearranged autosomes and long-term conservation of the X chromosome"

### Supplementary Figures

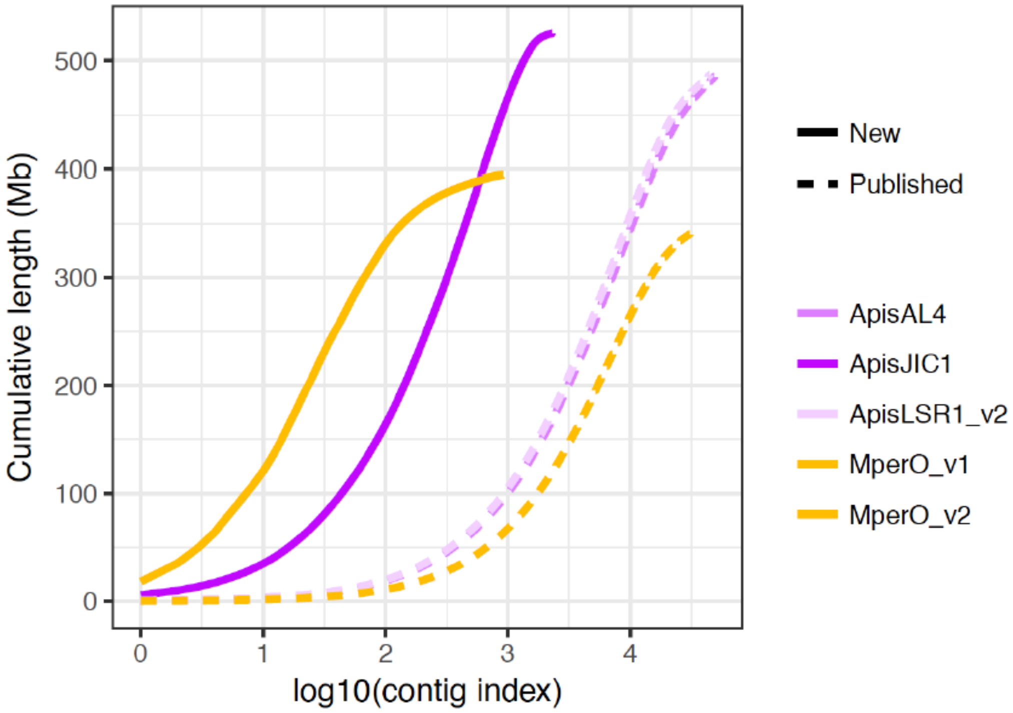


**Supplementary Figure 1:** Cumulative contig length for new a previously published assemblies of *Myzus persicae* and *Acyrthosiphon pisum* (see main text **Table 1** for details).

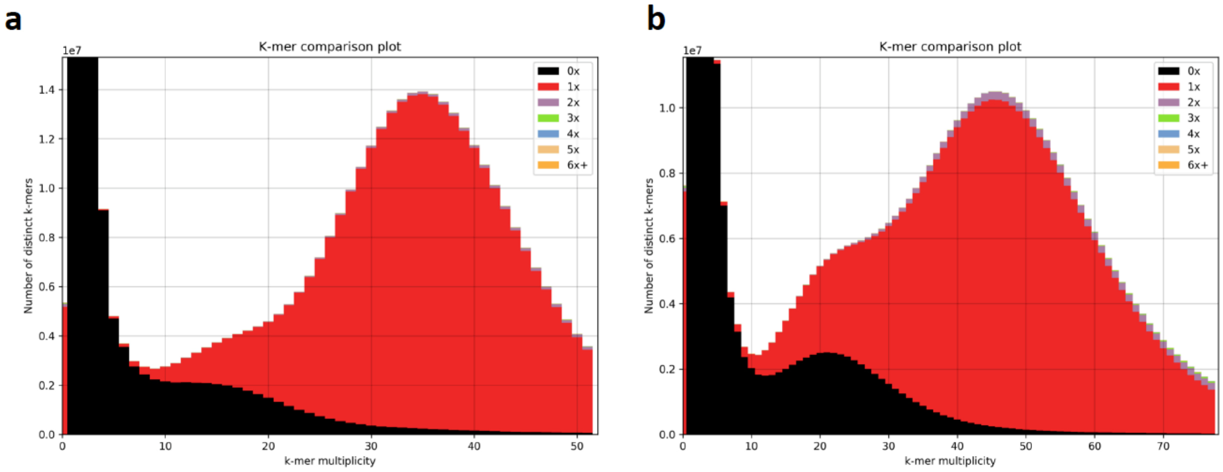


**Supplementary Figure 2:** (**a**) KAT (Mapleson et al. 2017) k-mer spectra plot comparing PCR free *M. persicae* clone O Illumina PE reads to MperO_v2 scaffolds. Colours indicate how many times fixed length words (k-mers) from the reads appear in the assembly. Red indicates k-mers found only once in the assembly, black indicates content present in the reads but missing from the assembly and other colours indicate k-mers that are duplicated in the assembly. The x-axis shows the number of times each k-mer is found in the reads (k-mer multiplicity) and the y-axis shows the count of distinct k-mers in 1x k-mer multiplicity bins. The bimodal distribution indicates heterozygosity in *M. persicae* clone O with the first peak at ~18x k-mer multiplicity corresponding to heterozygous genome content that has been collapsed in the assembly giving a haploid representation of the genome. The large peak at ~36x k-mer multiplicity corresponds to homozygous single copy genome content. The plot was generated with k=31. (**b**) As for (**a**) but comparing *A. pisum* JIC1 10x genomics linked reads (primers and barcodes removed) compared to ApisJIC1 scaffolds.

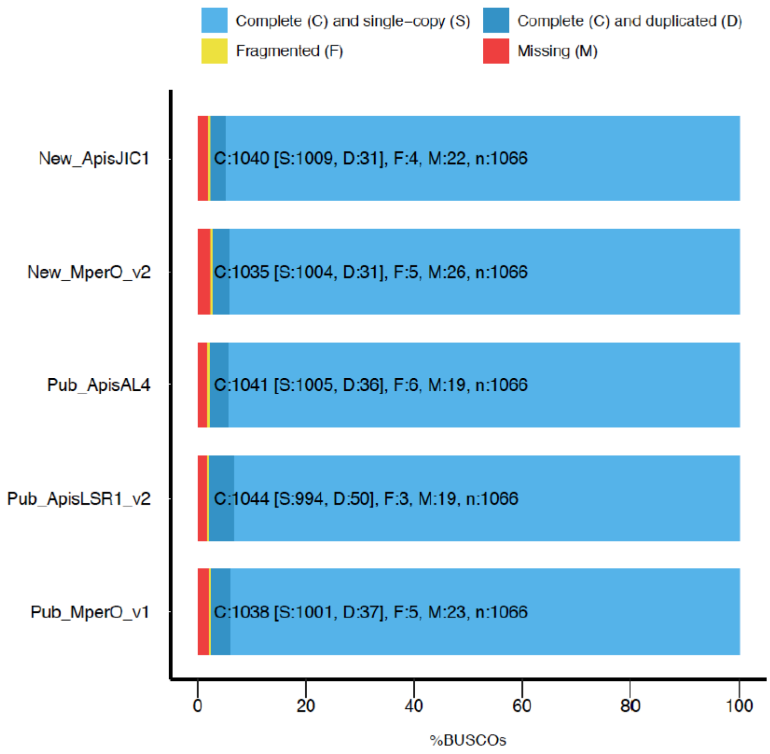


**Supplementary Figure 3:** BUSCO completeness plot for MperO_v2, ApisJIC1 and previously published genome assemblies of *A. pisum* (LSR1 v2 and Al4, see main text **Table 1** for details) and *M. persicae* clone O (v1.1). The genomes were assessed using the Arthropoda gene set (n=1,066).

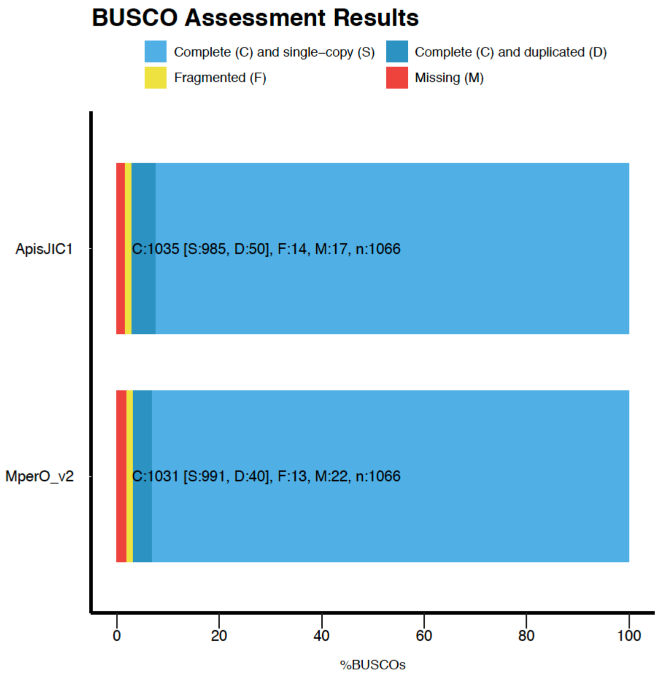


**Supplementary Figure 4:** BUSCO completeness plot for the BRAKER2 gene predications for MperO_v2 and ApisJIC1. BUSCO analysis was carried out using protein sequences with the longest transcript of each gene as the representative transcript and using the Arthropoda gene set (n=1,066).

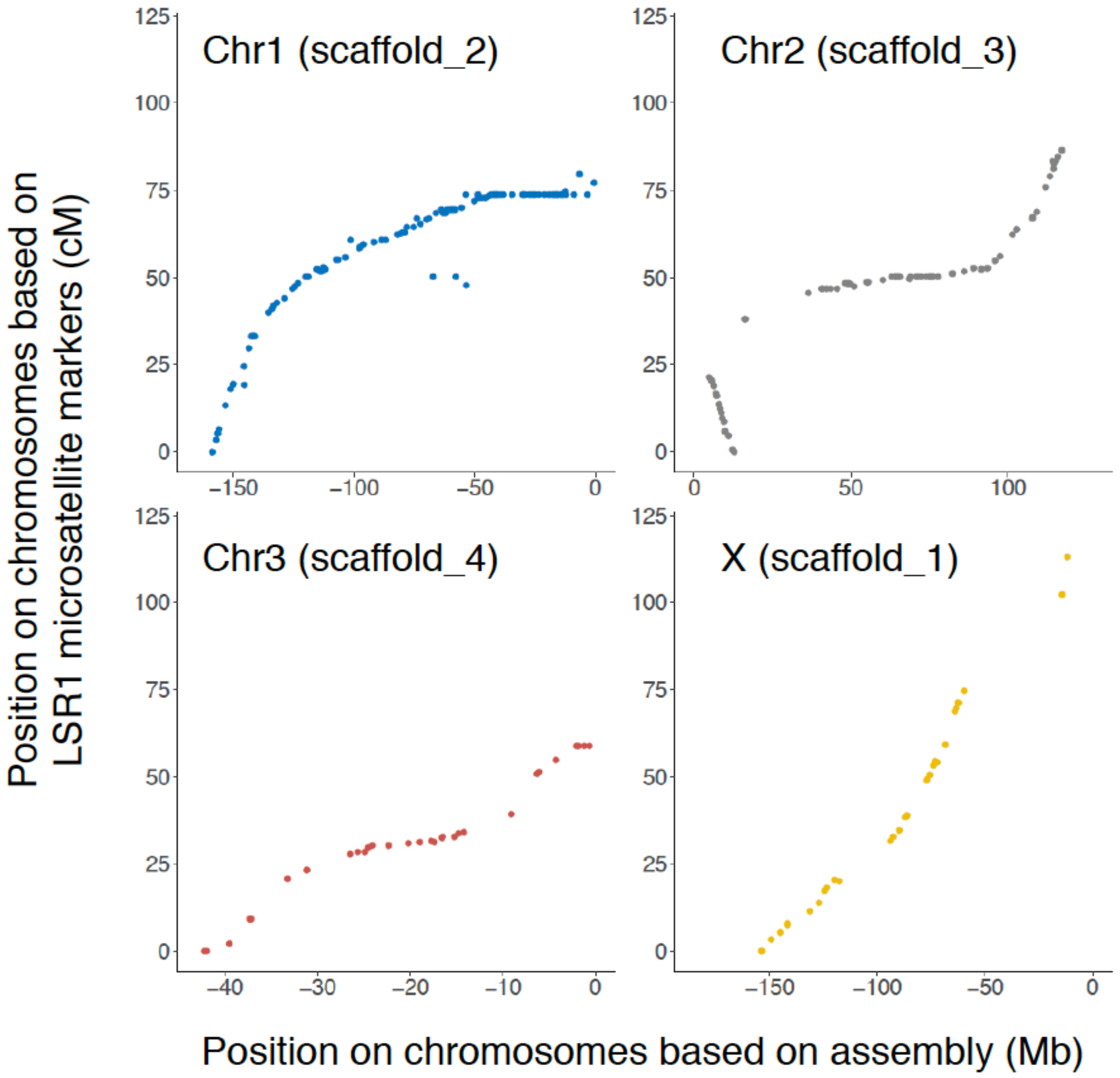


**Supplementary Figure 5:** Comparison of ApisJIC1 to the LSR1 linkage map (Jaquiéry *et al.* 2014). For each alignment the LSR1 chromosome ID is given with the ApisJIC1 scaffold ID in parenthesis. Mapping to the LSR1 linkage map was carried out as per Li et. al. (2019) with scripts from their github repository ([https://github.com/lyy005/Aphid_AL4_chromosome_assembly](https://github.com/lyy005/Aphid_AL4_chromosome_assembly)).

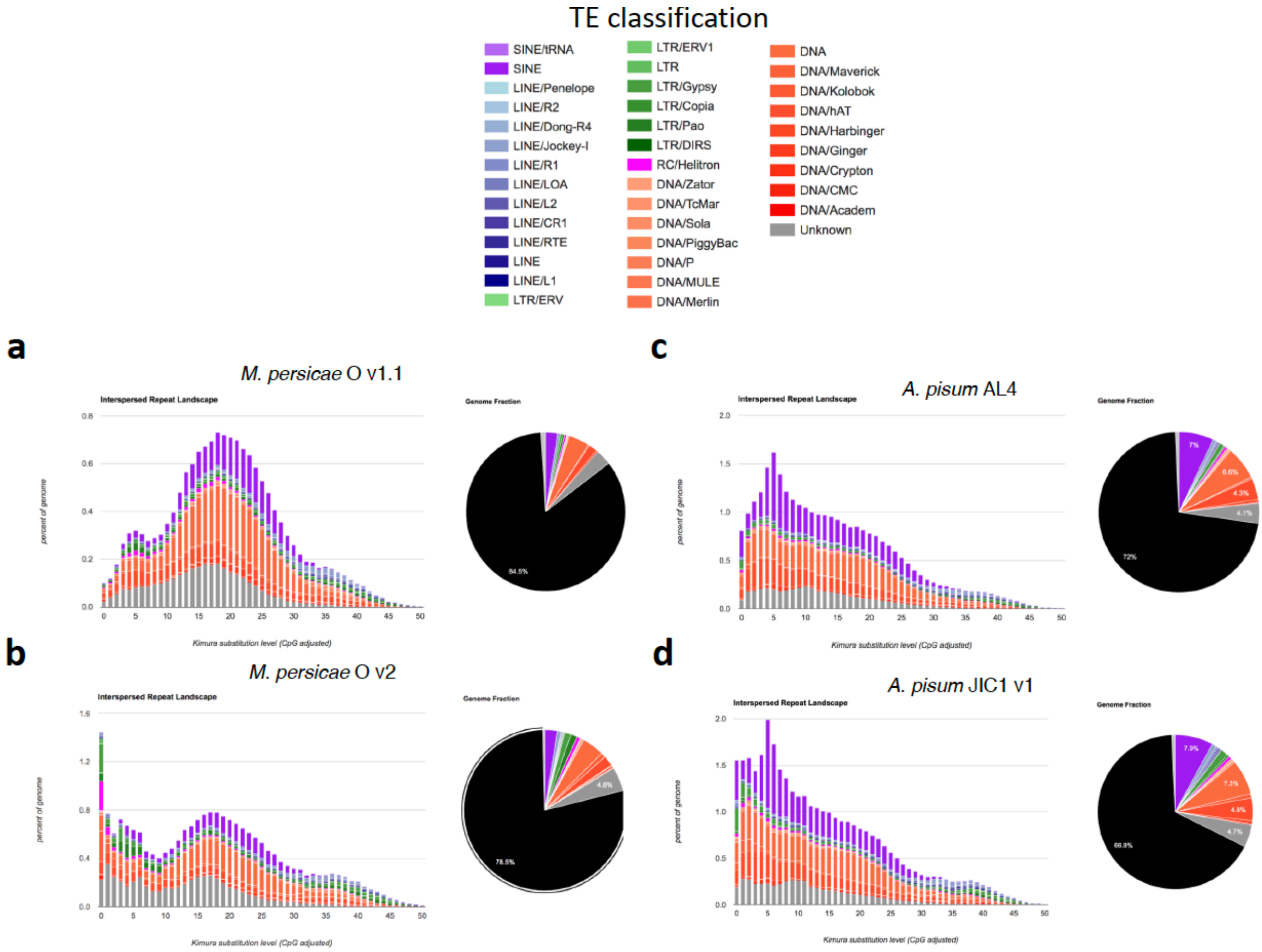


**Supplementary Figure 6:** Transposable element (TE) age distributions (*stacked histograms*, left) and genome proportions (*pie charts*, right) for previously published short-read and new long-read assemblies of *M. persicae* clone O and *A. pisum*. For *A. pisum*, short-read TE content is based on the AL4 assembly (Li et. al. 2019). Repeats were identified in each genome assembly using RepeatMasker. For each species, we used our newly generated de novo repeat library (RepeatModeler + RepBase insecta repeats; see Methods section). *Pie charts* show the percentage of bases in the assembly masked for each repeat type, with unmasked bases shown in black.

\

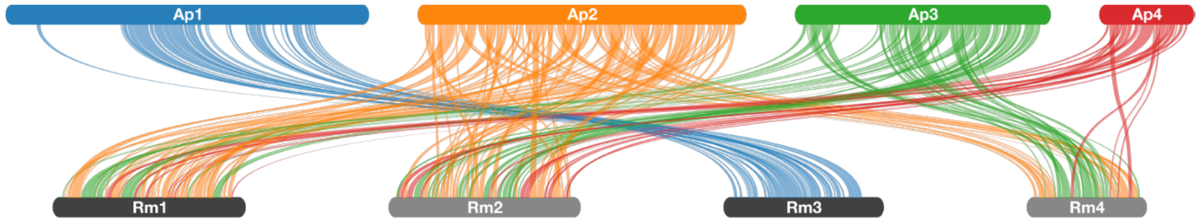


**Supplementary Figure 7:** Pairwise synteny relationships between A. pisum JIC1 and R. maidis. Links indicate the boundaries of syntenic blocks identified by MCscanX and are colour coded by *A. pisum* chromosome ID. Ap1 is the X chromosome.

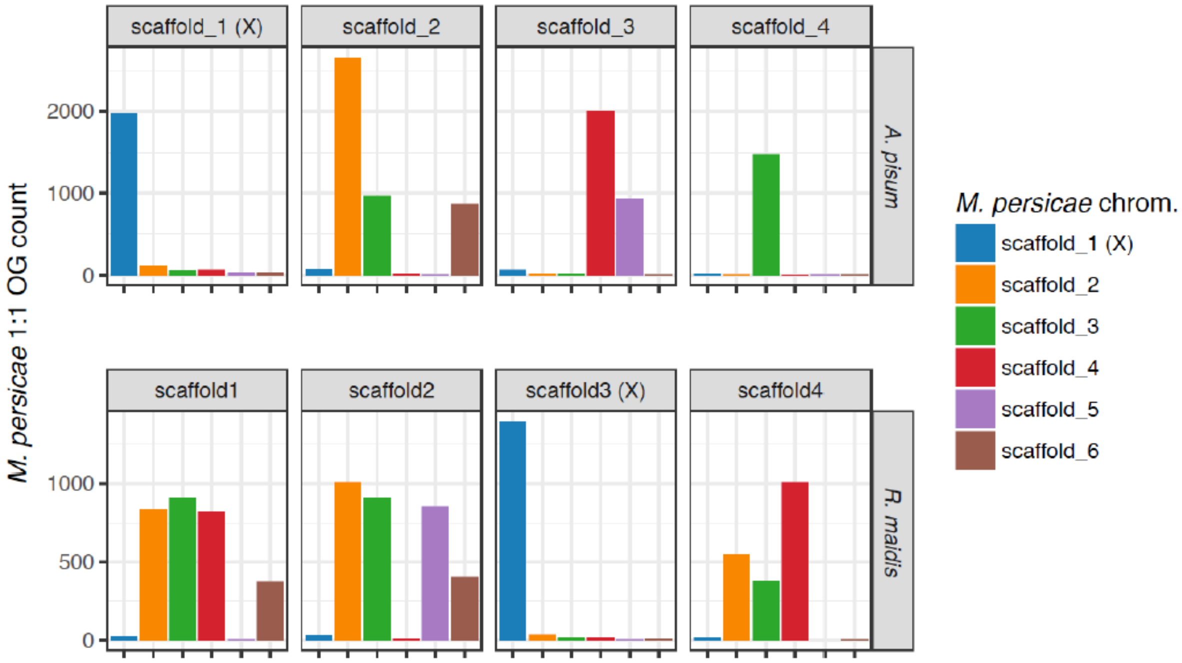


**Supplementary Figure 8:** Mapping of *M. persicae* clone O v2 one-to-one orthologs to *A. pisum* JIC1 (top; n= 11,372) and *R. maidis* (bottom; n = 9,594) chromosomes. One-to-one orthologs between *M. persicae* and each focal species were inferred based on gene trees with orthofinder. For each focal chromosome the count of orthologs from each of the six *M. persicae* chromosomes is given.

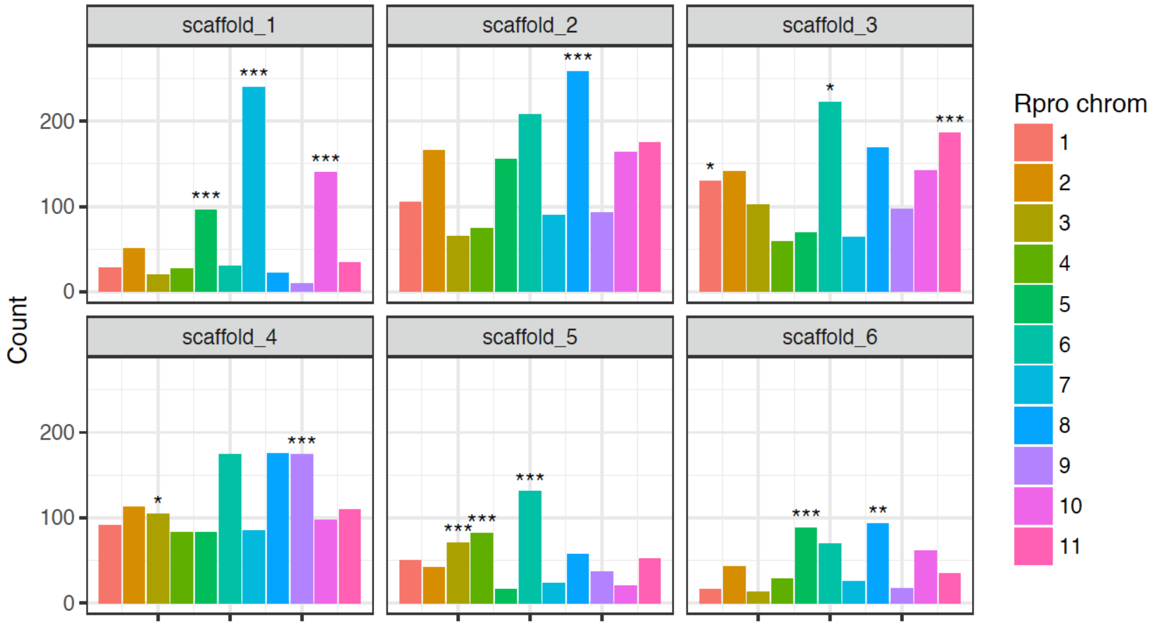


**Supplementary Figure 9:** Ortholog mapping between the aphid *Myzus persicae* and the kissing bug *Rhodnius prolixus*. Counts of *R. prolixus* chromosomal location for 5,992 *M. persicae* - *R. prolixus* 1:1 orthologs per *M. persicae* chromosome. Stars above bars indicate significant enrichment of a specific *R. prolixus* chromosome after correcting for multiple testing (BH corrected *binomial test*, *** = $p < 0.0001$, ** = $p < 0.001$, * = $p < 0.05$).

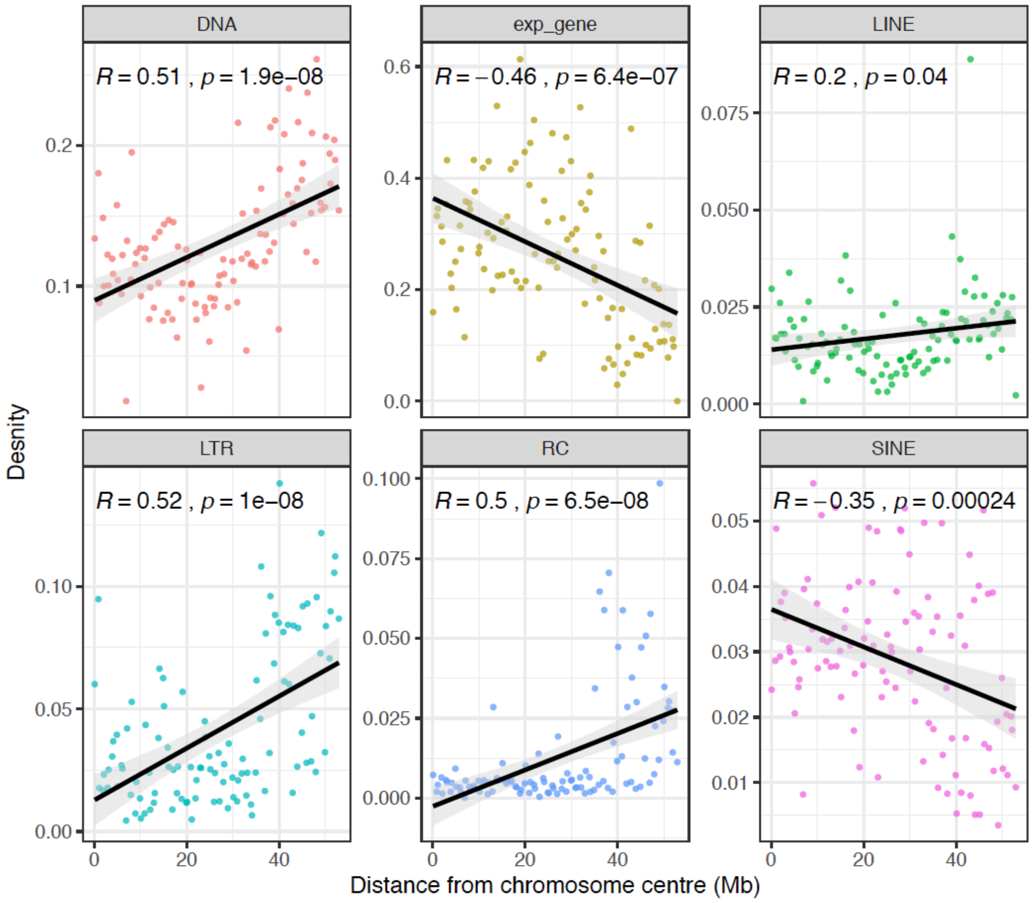


**Supplementary Figure 10:** *M. persicae* clone O v2 TE and expressed gene density vs distance from the centre of the X chromosome. Dots show feature density individual 1 Mb windows along the X chromosome. *Pearson correlation* (*R*) is reported for each feature.

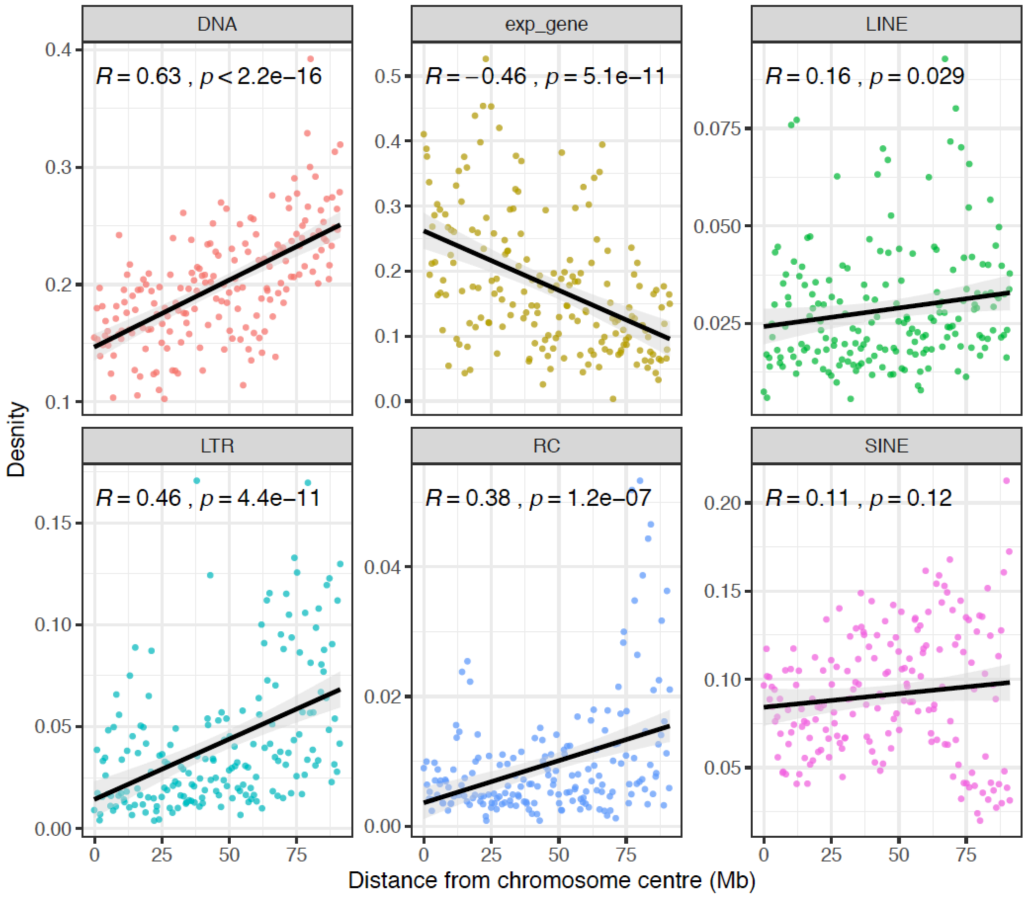


**Supplementary Figure 11:** *A. pisum* clone JIC1 v1 TE and expressed gene density vs distance from the centre of the X chromosome. Dots show feature density in individual 1 Mb windows along the X chromosome. *Pearson correlation* (*R*) is reported for each feature.

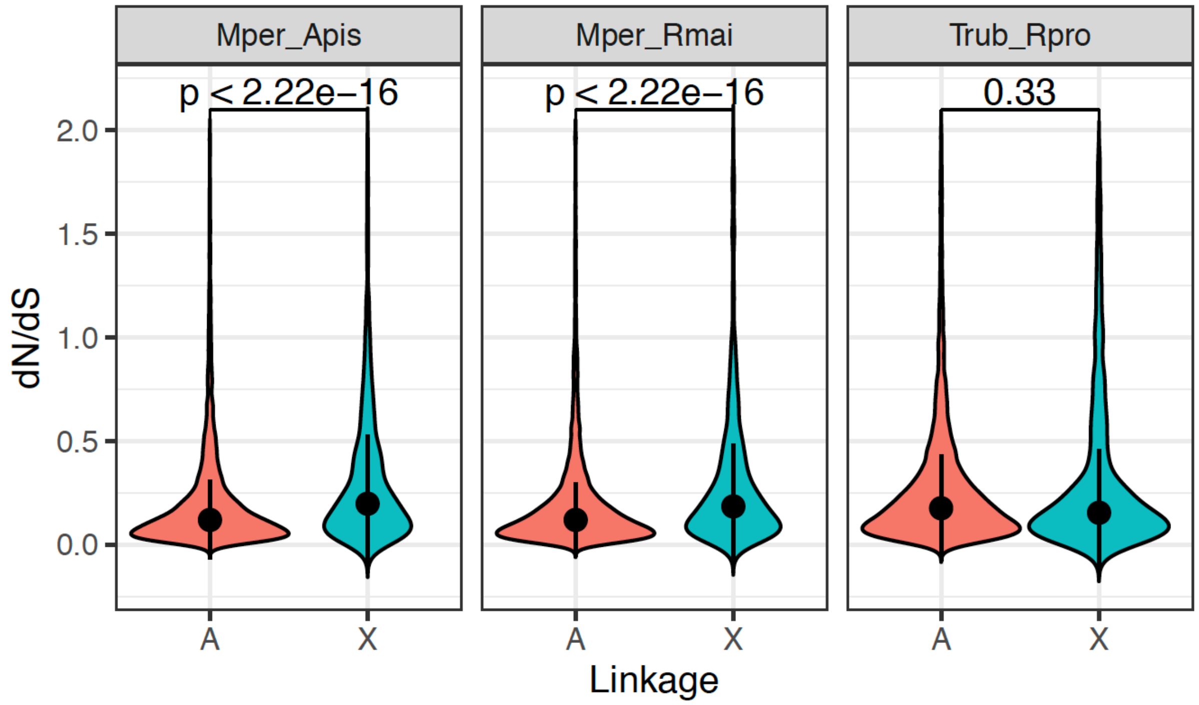


**Supplementary Figure 12:** Violin plots showing the rate of evolution ($d_N/d_S$) for X-linked (X) and autosomal (A) syntenic orthologs between *M. persicae* and *A. pisum* (Mper_Apis, n = 8,813), *M. persicae* and *R. maidis* (Mper_Rmai, n = 7,607) and *T. rubrofasciata* and *R. prolixus* (Trub_Rpro, n = 8,481). Ortholog pairs were retained that had $d_S > 0 < 2$ , $d_N > 0 < 2$ and $d_N/d_S < 2$. Black circles and whiskers show median and interquartile range, respectively. Numbers above comparisons show p values from *Wilcoxon rank-sum tests*. Orthologs were identified based on MCScanX synteny analysis.

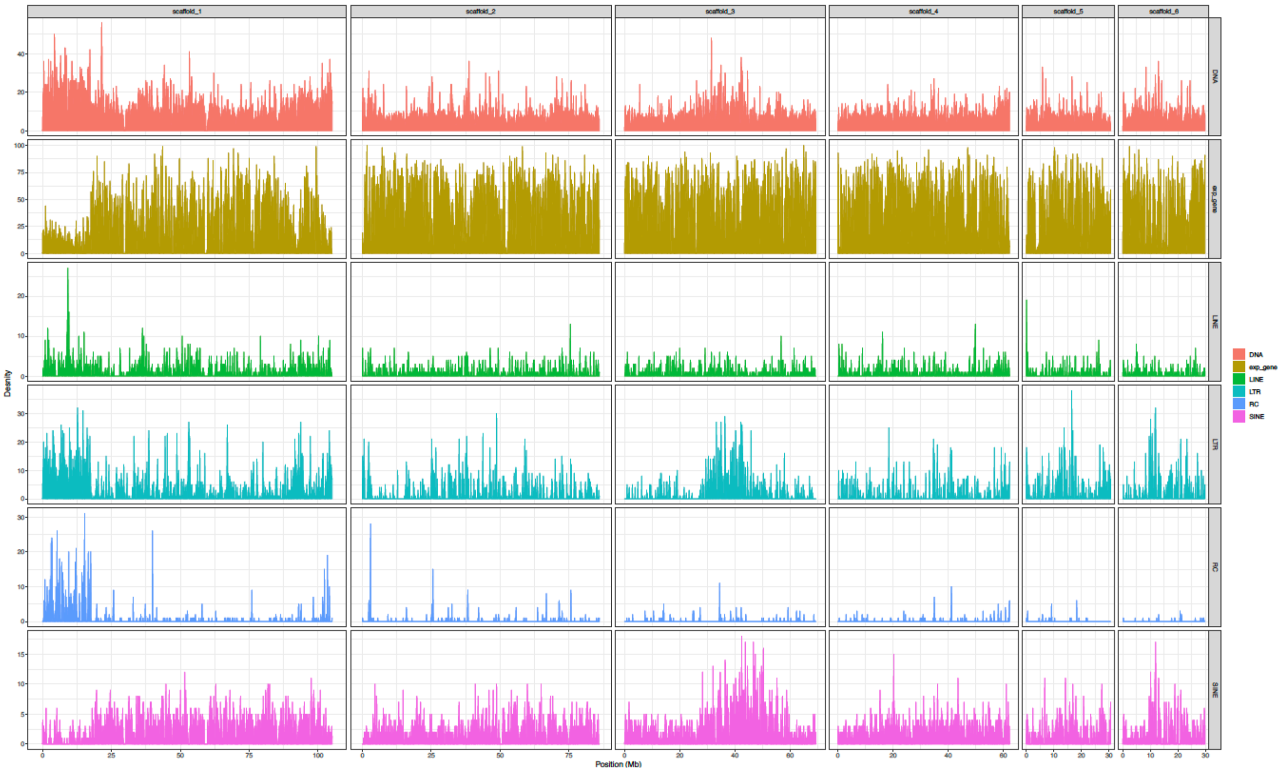


**Supplementary Figure 13:** The density of expressed genes (expr_gene) and transposable elements (DNA = DNA transposons, LINE = Long Interspersed Nuclear Elements, LTR = Long Terminal Repeat retrotransposons, RC = Rolling Circle transposons, SINE = Short Interspersed Nuclear Elements) across MperO_V2 chromosome-length scaffolds in 100 Kb fixed windows. Genes were classified as expressed if they had a Kallisto estimated read count > 4 in at least 12 / 24 *M. persicae* morph RNA-seq samples (see main text **Figure 5**).

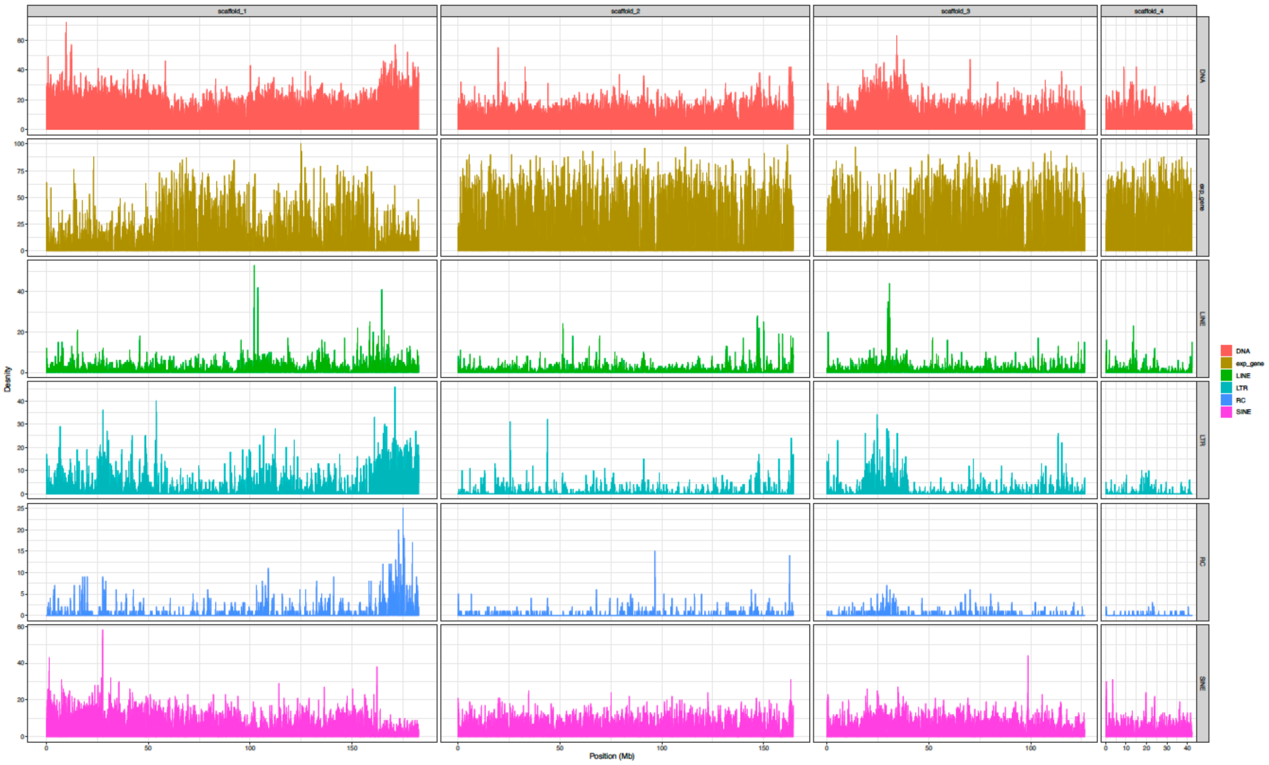


**Supplementary Figure 14:** As for (**Supplementary Figure 13**) but showing TEs and expressed genes across ApisJIC1 chromosome-length scaffolds. Genes were classified as expressed if they had a Kallisto estimated read count > 4 in at least 3 / 6 *A. pisum* morph RNA-seq samples from Jaquiéry et al. (2013).
