## Supplementary Note for "Chromosome-scale genome assemblies of aphids reveal extensively rearranged autosomes and long-term conservation of the X chromosome"

### Supplementary Note: *de novo* genome assembly of *M. persicae* clone O and *A. pisum* clone JIC1

#### *M. persicae* clone O

In a previous manuscript describing *M. persicae* genome sequence data (Mathers *et al.* 2017), a UK genotype (O) and a US genotype (G006) were sequenced and assembled using Illumina short-read technology. These genome assemblies have high gene-level accuracy but are fragmented, with 13,407 and 4,022 scaffolds, respectively (main text **Table 1**). Furthermore, both assemblies have a total length of ~355 Mb (including ~3% Ns) and are 54 Mb short of the 409 Mb genome size predicted for *M. persicae* by flow cytometry (Wenger *et al.* 2017). The missing content is most likely due to repetitive regions of the genome that are not well resolved by Illumina sequencing.

To create a new high-quality reference genome for *M. persicae* clone O, we sequenced 28 Gb (~70x coverage) of Nanopore long-reads that were used to generate a *de novo* assembly, and 24 Gb (~59x coverage) of 250 bp paired-end PCR free Illumina short-reads for polishing and quality control of the new assembly, and 123 Gb (~220x coverage) of *in vivo* HiC data for scaffolding (**Supplementary Table 6**).

##### *Assessing quality of preliminary contig assemblies*

To find out which assemblers work best for assembly of the *M. persicae* clone O Nanopore long-read data, we assessed the performance of three commonly used long-read *de novo* assemblers Canu v1.8 (Koren *et al.* 2017), Flye v2.4 (Kolmogorov *et al.* 2019) and wtdgb2 v2.3 (Ruan and Li 2019). Assemblies with canu and Flye were generated using default Nanopore parameters ("--nano-raw" and "-nanopore-raw", respectively). For wtdgb2 we used default Nanopore settings ("-x ont") and various values of the "-p" and "-k" parameter on a subset of reads at least 15Kb long (17.5 Gb, ~43x coverage) were used (**Table 1**). Each assembly was polished using three rounds of Pilon v1.22 (Walker *et al.* 2014) and the quality of the assemblies was assessed based on contiguity, completeness and the amount of pseudo duplication caused by the assembly of haplotigs (heterozygous regions assembled as separate contigs). Contiguity statistics were gathered with abyss-fac (Simpson *et al.* 2009; Jackman *et al.* 2017). Completeness and the level of duplication in the assemblies was assessed by comparing k-mers in our PCR-free Illumina reads to k-mers in each candidate assembly with KAT comp from the k-mer analysis toolkit (KAT) v2.3.4 (Mapleson *et al.* 2017) and by identifying conserved genes with BUSCO v3 (Simão *et al.* 2015; Waterhouse *et al.* 2018) using the Arthropoda gene set (n=1,066).

**Table 1:** Statistics for Nanopore assemblies generated with Canu, Flye and wtdbg2.

| Assembly | Size (Mb) | N50 (Mb) | Longest contig (Mb) |
|---|---|---|---|
| canu_default | 534.8 | 3.07 | 22.66 |
| Flye_default | 439.8 | 1.31 | 6.83 |
| wtdbg2_ont_default | 402.6 | 1.23 | 13.39 |
| wtdbg2_p0_k15_L15Kb | 403 | 1.71 | 18.51 |
| wtdgb2_p0_k17_L15Kb | 412 | 2.12 | 14.99 |
| wtdbg2_p0_k19_L15Kb | 410.6 | 2.29 | 12.36 |
| wtdgb2_p19_k0_L15Kb | 401.7 | 1.83 | 23.08 |

All the candidate assemblies had low levels of missing content after polishing with pilon (**Figure 1**). The canu assembly had the highest contiguity (**Table 1**; N50 = 3.07 Mb), though the assembly size was much larger than the predicted genome size (535 Mb vs 409 Mb). K-mer analysis showed that the canu assembly suffered from high levels of duplication as indicated by large numbers of k-mers found twice in the assembly but for which coverage in the raw reads was similar to that of single-copy content (i.e. k-mers should be found only once in a haploid assembly) (**Figure 1a**). BUSCO analysis also shows high duplication levels in the canu assembly (13.7% duplicated). The Flye assembly was also larger than the predicted genome size (439 Mb vs 409 Mb) and had moderate levels of duplication according to BUSCO (4.3% duplicated) and k-mer analysis (**Figure 1b**). However, contiguity of the Flye assembly was approximately half that that of the canu assembly (1.31 Mb vs 3.07 Mb) and similar to the wtdgb2 assembly with default Nanopore parameters (1.23 Mb). Modifying the default wtdbg2 parameters resulted in an almost doubling of contiguity (2.29 Mb vs 1.23 Mb) and a larger assembly size (412 Mb vs 403 Mb) for the largest wtdgb2 ("-p 0 -k 17 -L 15000") assembly (**Table 1**). The wtdgb2_p0_k17_L15kb assembly was highly complete and had the lowest levels of duplication based on the K-mer and BUSCO analyses (**Figure 1c**).

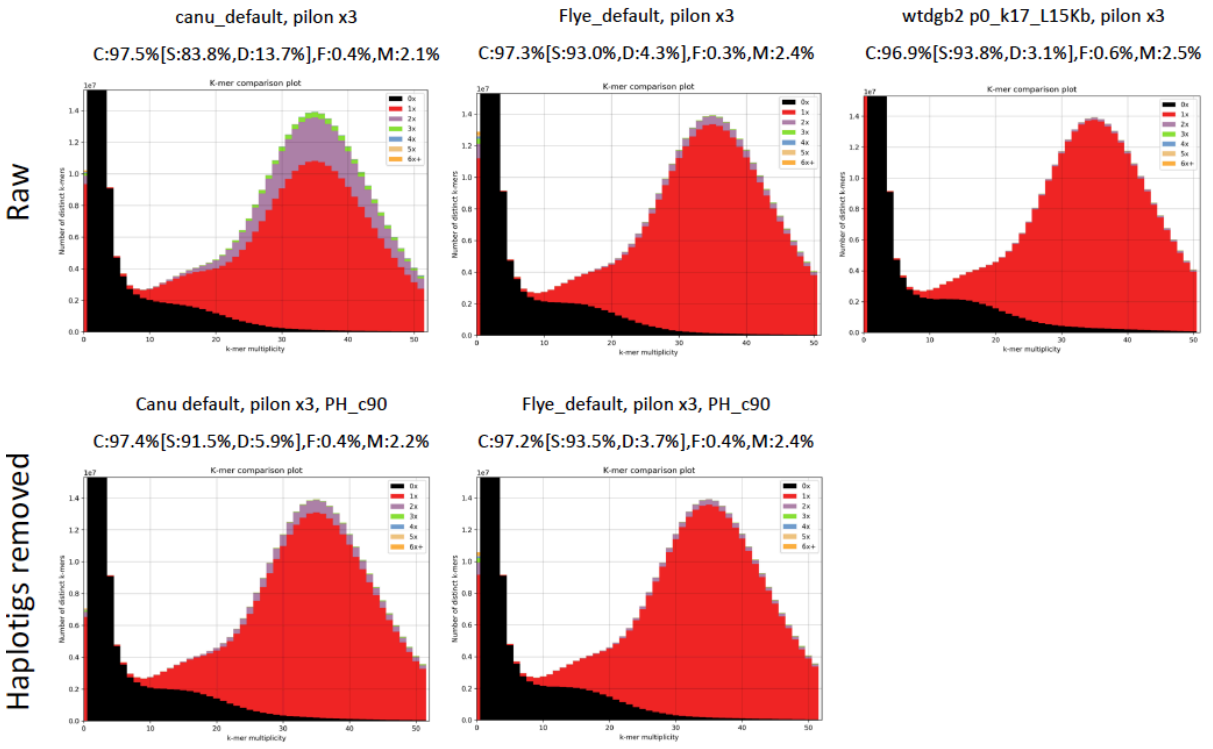


**Figure 1:** KAT k-mer spectra plots comparing PCR free *M. persicae* clone O Illumina PE reads to long-read assemblies generated with canu and Flye with default parameters and wtdgb2 with the parameters “-x ONT -p 0 -k 17 -L 15000”. Bottom row shows the same assemblies after removal of putative haplotigs with purge_haplotigs. Each assembly was polished with three rounds of short-read polishing with Pilon. Colours indicate how many times fixed length words (k-mers) from the reads appear in the assembly. Red indicates k-mers found only once in the assembly, black indicates content present in the reads but missing from the assembly and other colours indicate k-mers that are duplicated in the assembly. The x-axis shows the number of times each k-mer is found in the reads (k-mer multiplicity) and the y-axis shows the count of distinct k-mers in 1x k-mer multiplicity bins. The bimodal distribution indicates heterozygosity in *M. persicae* clone O with the first peak at ~18x k-mer multiplicity corresponding to heterozygous genome content that has been collapsed in the assembly giving a haploid representation of the genome. The large peak at ~36x k-mer multiplicity corresponds to homozygous single copy genome content. The plot was generated with k=31. BUSCO scores for each assembly expressed as the percentage of BUSCO Arthropoda genes (n = 1,066) found in each category are given above the K-mer spectra: C = complete, S = single-copy and complete, D = complete and duplicated, F = fragmented, M = missing.

Given the high levels of duplication found in our canu and Flye assemblies we attempted to remove contigs likely to be haplotigs from each assembly. Various tools have been developed to remove haplotigs from genome assemblies (Pryszcz and Gabaldón 2016; Roach *et al.* 2018). We tested purge_haplotigs (Roach *et al.* 2018), which uses a combination of whole genome self-alignment and coverage patterns (based on long-read alignment), to identify candidate haplotigs and remove them from the draft assembly. We compared BUSCO and KAT comp runs before and after purge_haplotigs to investigate how effectively duplicated content is removed from each assembly and to test whether purge_haplotigs erroneously removes genuine single-copy genome content. To set the coverage bounds required by purge_haplotigs we aligned our Nanopore data to the canu assembly with minimap2 v2.14-

r883 (Li 2018) using default Nanopore parameters ("-ax map-ont") and plotted a coverage histogram (**Figure 2**). Based on visual inspection of the histogram, we set upper coverage thresholds to assign contigs as either low coverage contamination ("-l 9"), candidate haplotigs ("-m 45") and high coverage repeats ("-h 92"). Candidate haplotigs were removed from the assembly covered if the covered at least 90% of another, longer, contig in the assembly. Purge_haplotigs reduced the amount of duplicated content in both the canu (**Figure 1d**) and Flye (**Figure 1e**) assemblies without reducing genuine single copy content. However, even after deduplication, both the canu and flye assemblies still had a greater amount of duplicated content than the wtdgb2 assembly (**Figure 1c-e**). We therefore focused on further improving the wtdgb2 contig assembly (wtdgb2_p0_k17_L15Kb).

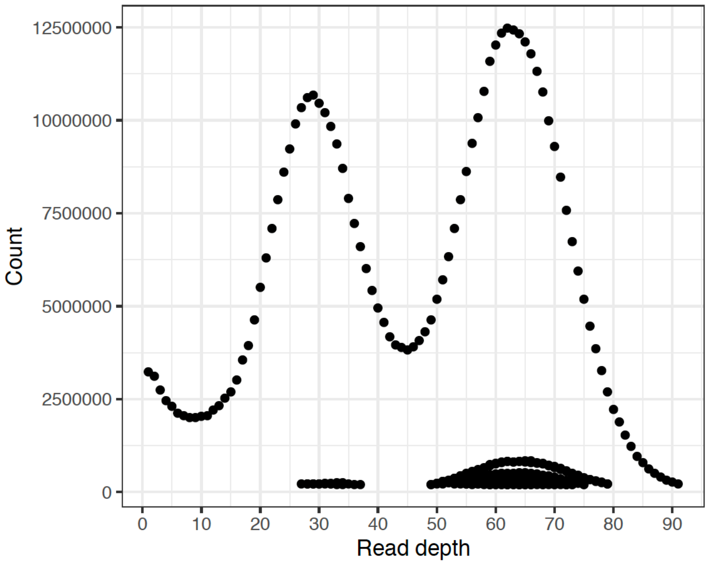


**Figure 2:** purge_haplotigs coverage histogram for *M. persicae* clone O Nanopore reads aligned to the canu assembly.

##### ***Final contig assembly***

Assemblies generated with different software or parameters may contain complementary contiguity and genome content (Chakraborty *et al.* 2016). To investigate whether this was the case for our wtdgb2 assemblies, we generated a whole genome alignment between wtdgb2_p0_k17_L15Kb (the wtdgb2 assembly with the most assembled content) and wtdgb2_p19_k0_L15Kb (the wtdgb2 assembly with the longest contig) using mashmap v2 (Jain *et al.* 2018) (**Figure 3a**). This showed extensive complementary contiguity between the two assemblies. We also found non-overlapping sets of complete BUSCO genes in the two assemblies (**Figure 3b**). Therefore, quickmerge v0.3 (Chakraborty *et al.* 2016) was used to combine the two assemblies into a final contig assembly.

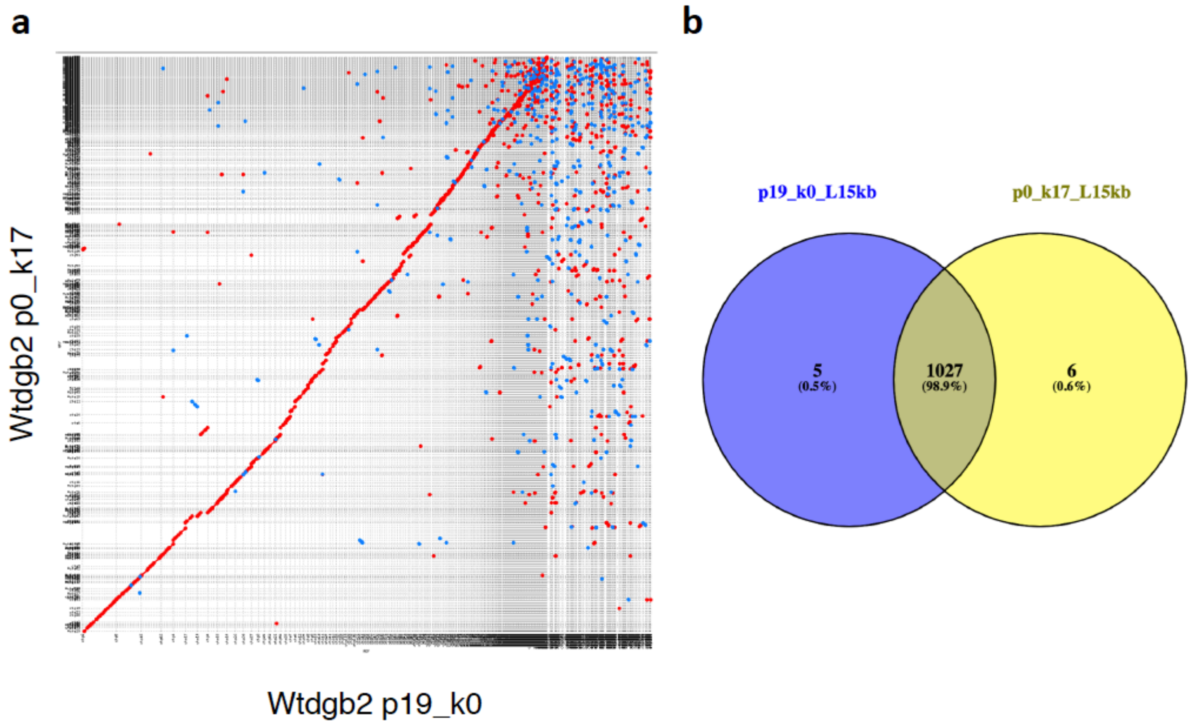


**Figure 3:** Complementary contiguity between *M. persicae* clone O wtdgb2 assemblies with different parameters. (**a**) *Dot plot* showing a mashmap whole genome alignment between a wtdgb2_p19_k0_L15kb (reference, x-axis) and wtdgb2_p0_k17_L15kb (query, y-axis). (**b**) *Venn diagram* showing overlap of complete BUSCO Arthropoda genes found in the two assemblies.

Although we carried out assembly evaluation in the previous sections using Pilon polished assemblies (3 rounds short-read polishing) we wanted to include long-read polishing for our final assembly. Therefore, before running quickmerge, we polished each raw wtdgb2 input assembly with the wtdgb2 polisher (wtpoa-cns) using the full set of Nanopore reads aligned with minimap2. We then carried out additional polishing using long- and short-reads after assembly merging (see section below). quickmerge requires the designation of one assembly as the *reference* and one assembly as the *query* and uses the *reference* sequence to join contigs in the *query* sequence (https://github.com/mahulchak/quickmerge/wiki). As such, the resulting merged assembly most closely resembles the content of the *query* assembly. Therefore, wtdgb2_p0_k17_L15Kb was set as the *query* (as it has the most assembled content) and wtdgb2_p19_k0_L15Kb as the *reference*, and merged the assemblies using quickmerge parameters as per recommendations in the documentation (https://github.com/mahulchak/quickmerge). Specifically, the length cut-off for anchor contigs ("-l") was set to the N50 of the *reference* assembly (1.83 Mb; **Table 2**), and the minimum alignment length was considered for merging ("-lm") to 10 Kb. This produced a merged assembly that was more complete and more contiguous than either input assembly (**Table 2**).

**Table 2:** Assembly statistics for wtdgb2 assemblies before and after merging with quickmerge and after polishing ("_polished" = three rounds of racon and three rounds of Pilon) and haplotig removal ("_PH_c90" = purge_haplotigs filtered assembly using 90% alignment coverage threshold).

| Assembly | Size (Mb) | N50 (Mb) | Longest contig (Mb) |
|---|---|---|---|
| wtdgb2_p0_k17_L15Kb | 405.9 | 2.11 | 14.99 |
| wtdgb2_p19_k0_L15Kb | 396.4 | 1.85 | 22.99 |
| merged | 408.2 | 4.38 | 24.35 |
| merged_polished | 412.6 | 4.42 | 24.56 |
| merged_polished_PH_c90 | 397.7 | 5.09 | 24.56 |

***Assembly polishing***

To increase base-level accuracy of the merged assembly three rounds of long-read polishing with racon v1.3.1 (Vaser *et al.* 2017) were conducted, followed by three rounds of short-read polishing with Pilon using our PCR free Illumina data. The effect of both long- and short-read polishing with KAT comp and BUSCO was assessed after each iteration (**Figure 4**). The polished assembly contained a small amount of duplicated content (**Figure 4**), which was removed using purge_haplotigs with parameters as described above for the canu and Flye test assemblies. The final assembly is contiguous (N50 = 5.09 Mb, longest contig = 24.56 Mb; **Table 2**), has low levels of duplication and is highly complete (**Figure 4d**).

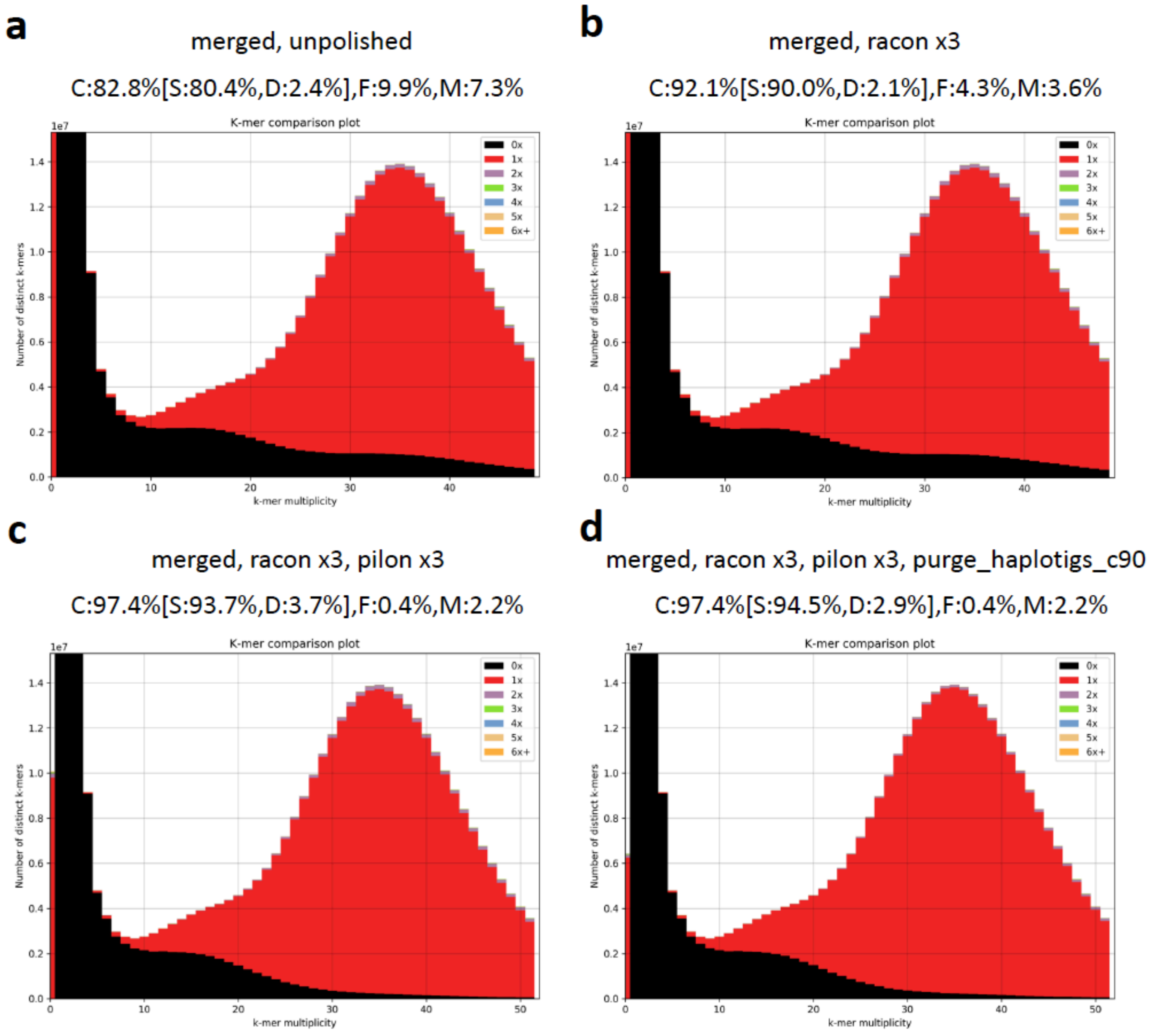


**Figure 4:** KAT k-mer spectra plots comparing PCR free *M. persicae* clone O Illumina PE reads to: the unpolished merged wtdgb2 assembly (**a**) and the same assembly after long-read polishing with three iterations of racon (**b**), three iterations of racon and three iterations of Pilon short-read polishing (**c**) and after removal of putative haplotigs from the polished assembly (**d**). BUSCO scores are shown above each K-mer spectra as per Figure 1.

##### *HiC scaffolding*

HiC scaffolding of the final contig assembly was carried out as described in the main text using Juicer v1.6.2 (Durand *et al.* 2016) to identify HiC contacts and the 3D-DNA assembly pipeline (Dudchenko *et al.* 2017) to correct misassemblies in the input contig assembly and then to order contigs (or scaffolds for *A. pisum* JIC1) into super-scaffolds. Further manual corrections of the initial 3D-DNA assembly were performed with Juicebox Assembly Tools (JBAT). The initial 3D-DNA assembly is shown in **Figure 6a** and the reviewed assembly, after assembly errors have been corrected, is shown in **Figure 2b**. Following manual review, genomic content removed from super-scaffolds by false positive manual edits was reintegrated with the 3D-DNA module seal to create a final scaffolded assembly.

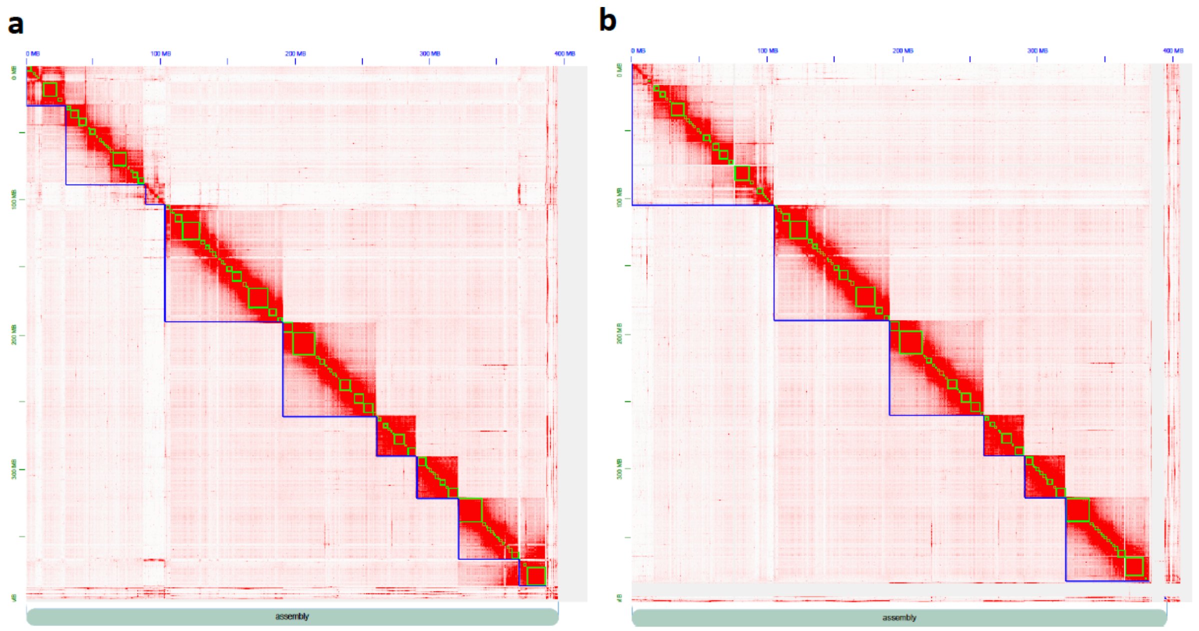


**Figure 6:** Heatmaps showing frequency of HiC contacts along the 3d-DNA scaffolded *M. persicae* clone O contig assembly before (**a**) and after (**b**) manual review with JBAT. Blue lines indicate super scaffolds and green lines show contigs. The X and Y axis show cumulative length in Mb.

##### *Contamination filtering and final quality control*

We checked the final scaffolded assembly for contamination with BlobTools (Kumar *et al.* 2013; Laetsch and Blaxter 2017). BlobTools generates taxon annotated GC content-coverage plots of the average GC content and read coverage for each scaffold in an assembly, and assigns taxonomy to each scaffold based on blast searches to the NCBI database. Each scaffold in the HIC assembly was annotated with taxonomy information based on blastn v2.2.31 (Camacho *et al.* 2009) searches against the NCBI nucleotide database (nt, downloaded 13/10/2017) with the options "-outfmt '6 qseqid staxids bitscore std sscinames sskingdoms stitle' -culling_limit 5 - evalue 1e-25". Two runs of BlobTools were carried out, one using coverage estimated from the PCR free Illumina short-reads and a second run using coverage estimated from the Nanopore long-reads. For the short-read run, PCR-free Illumina reads were mapped to the HiC assembly with BWA-MEM v0.7.7 (Li 2013) using default settings. The resulting BAM file was sorted with SAMtools v1.3 (Li *et al.* 2009) and passed to BlobTools along with the table of blastn results. For the long-read run, we mapped the full set of long-reads to the HiC assembly with minimap2 v2.14-r883 (Li 2018) using default Nanopore parameters ("-ax map-ont") and post possessed the alignments as per the Illumina short reads. Manual inspection of the taxon annotated GC content-coverage plots revealed low levels of contamination (**Figure 7a** and **b**). To make the final assembly, scaffolds assigned to the aphid *Buchnera aphidicola* endosymbiont at the genus-level (n = 82, total content = 623,074 Kb) were removed from the assembly. Scaffolds that likely correspond to low coverage contamination based on the criteria of having less than 10x average coverage in the Illumina reads and the Nanopore long-reads (n = 133, total content = 1.13 Mb) were also removed from the assembly. In the final *M. persicae* clone O v2 release, remaining scaffolds were renamed and ordered by size with SeqKit v0.9.1 (Shen *et al.* 2016). The assembly was then checked a final time with KAT comp and BUSCO (**Supplementary Figures 2** and **3**).

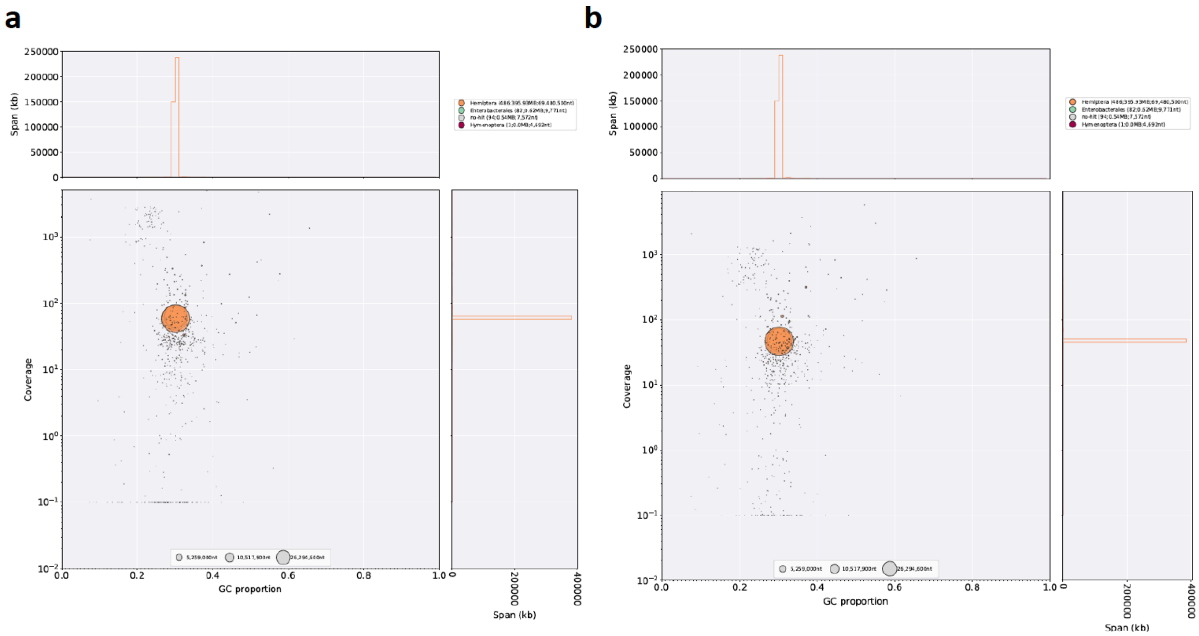


**Figure 7:** Taxon-annotated GC content-coverage plots of the *M. persicae* clone O HiC scaffolded assembly. Each circle represents a scaffold in the assembly, scaled by length, and coloured by order-level NCBI taxonomy assigned by BlobTools. The X axis corresponds to the average GC content of each scaffold and the Y axis corresponds to the average coverage based on alignment (**a**) Nanopore long-reads or (b) PCR free Illumina short reads. Marginal histograms show cumulative genome content (in Kb) for bins of coverage (Y axis) and GC content (X axis).

#### *A. pisum* JIC1

To create a new high-quality reference genome for *A. pisum* clone JIC1, we sequenced 18 Gb (~35x coverage of the *A. pisum* genome) of Nanopore long-reads for *de novo* assembly, 45 Gb of 150 bp 10X Genomics linked-reads (~88x coverage) to scaffold and polish the long-read contig assembly, and 21 Gb (~40x coverage) of *in vivo* HiC data for further scaffolding into chromosome length super scaffolds (**Supplementary Table 6**).

##### *Contig assembly*

As for *M. persicae* clone O, we generated multiple long-read contig assemblies to identify the most effective assembler for our data. Due to its poor performance and long run time with *M. persicae* clone O, assembly of the *A. pisum* JIC1 Nanopore data with canu was not attempted. Instead, assemblies were generated with Fly using Nanopore defaults and wtdgb2 using two sets of parameters that gave the best results for the *M. persicae* data: "-x ONT -p 0 -k 17 -L15000" and "-x ONT -p 19 -k 0 -L15000". Each assembly was polished with 3 rounds of racon using all long reads and 3 rounds of pilon using 10x Genomics linked reads stripped of their primers and barcodes. Assembly quality was assessed using BUSCO and KAT comp. The processed 10X Genomics linked reads were used to generate k-mer spectra with KAT comp.

The most contiguous assemblies were generated with wtdgb2 (Table 3). Both wtdgb2 parameter sets generated assemblies with contig N50s > 1.5 Mb and contigs over 20 Mb in length. However, k-mer analysis revealed that the wtdgb2 assemblies had extensive missing content (**Figure 7b** and **c**). In contrast, Flye produced a more fragmented assembly (contig N50 = 0.5 Mb, longest contig 5.79 Mb). However, the Flye assembly contained nearly 80 Mb more content than either of the wtdgb2 assemblies. The increase in content is most likely due to both reduced missing content and increased duplication, caused by the assembly of haplotigs (**Figure 7a**).

**Table 3:** Statistics for *A. pisum* JIC1 Nanopore assemblies generated with Flye and wtdbg2.

| Assembly | Base pairs (Mb) | Contig N50 (Mb) | Longest contig (Mb) |
|---|---|---|---|
| flye_default | 581.1 | 0.5 | 5.79 |
| wtdgb2_p0_k17_L15Kb | 504.6 | 1.62 | 22.15 |
| wtdgb2_p19_k0_L15Kb | 501.8 | 1.73 | 20.9 |

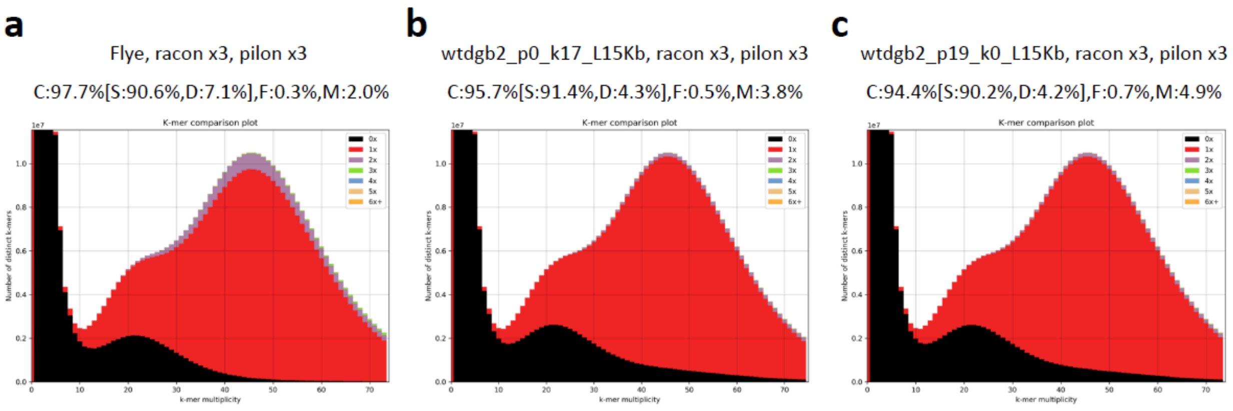


**Figure 7:** KAT k-mer spectra plots comparing *A. pisum* JIC1 10X Genomics reads to long-read assemblies generated with (**a**) Flye, (**b**) wtdgb2 with the parameters "-x ONT -p 0 -k 17 -L 15000" and (**c**) wtdgb2 with the parameters "-x ONT -p 19 -k 0 -L 15000". BUSCO scores are shown above each k-mer spectra. Each assembly was polished three times with racon and three times with Pilon prior to running QC.

Based on these preliminary results, two strategies to generate a contiguous and complete assembly were attempted. First, the two wtgdbg2 assemblies were merged with quickmerge using the parameters "-l 1598215 -ml 10000" and setting the more complete wtdgb2_p0_k17_L15Kb assembly as the query. This produced an assembly that was highly contiguous (N50 = 2.76 Mb, longest contig 29.02 Mb), but contained more missing content than the Flye assembly (**Figure 8a; Table 4**). Next, to increase the contiguity of the highly complete Flye assembly, this assembly and the wtdgb2_p0_k17_L15Kb assembly were merged with quickmerge, using the Flye assembly as the query and the parameters "-l 505810 -ml 10000". This dramatically increased contiguity (N50 = 3.02 Mb, longest contig 28.53 Mb). However, k-mer analysis showed that although the assembly is complete it still contained a high level of duplication (**Figure 8b; Table 4**).

**Table 4:** Statistics for merged *A. pisum* JIC1 Nanopore assemblies generated with quickmerge.

| Assembly | Base pairs (Mb) | Contig N50 (Mb) | Longest contig (Mb) |
|---|---|---|---|
| merged_wtdgb2 | 507.60 | 2.76 | 29.02 |
| merged_Flye_wtdgb2 | 571.00 | 3.02 | 28.53 |

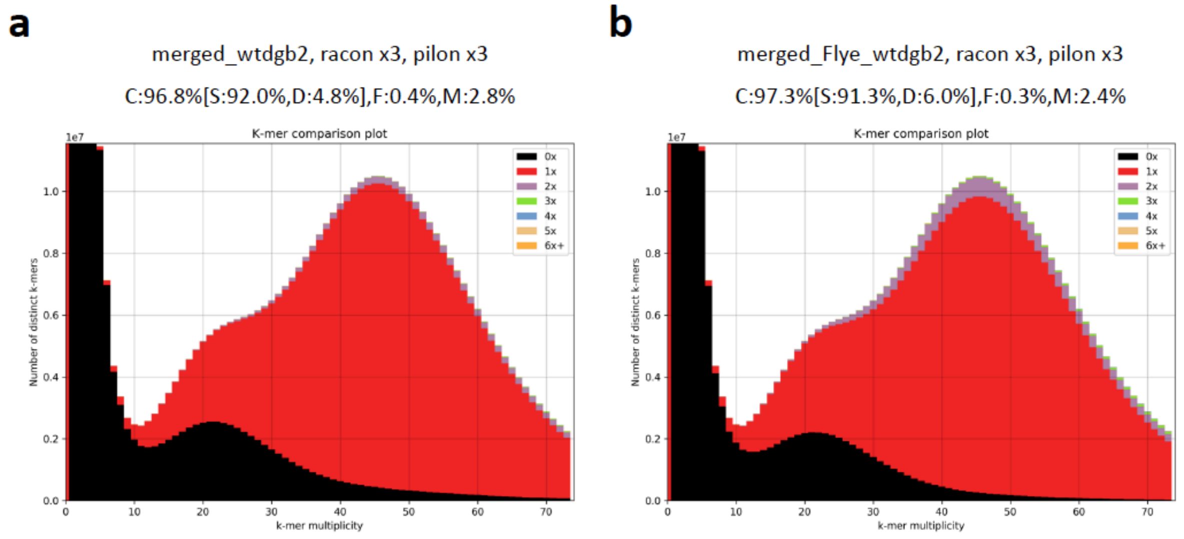


**Figure 8:** KAT k-mer spectra plots for *A. pisum* JIC1 long-read assemblies merged with quickmerge. (**a**) quickmerge assembly generated from merging wtdgb2_p0_k17_L15Kb and wtdgb2_p19_k0_L15Kb. (**b**) quickmerge assembly generated from merging Flye and wtdgb2_p0_k17_L15Kb assemblies. BUSCO scores are shown above each K-mer spectra.

To reduce duplication levels in the merged Flye-wtdgb2 assembly, the assembly was filtered with purge_haplotigs. We aligned our Nanopore data to the Flye assembly with minimap2 using default Nanopore parameters ("-ax map-ont") and plotted a coverage histogram (**Figure 9**). Based on this histogram, we set upper coverage thresholds to assign contigs as either low coverage contamination ("-l 4"), candidate haplotigs ("-m 21") and high coverage repeats ("-h 57"). We ran purge_haplotigs twice, once using a coverage threshold of 90% and once using a more relaxed setting of 75% (**Figure 10a** and **b**). Both runs reduced duplication levels compared to the unfiltered assembly, though k-mer analysis showed that residual duplication remained.

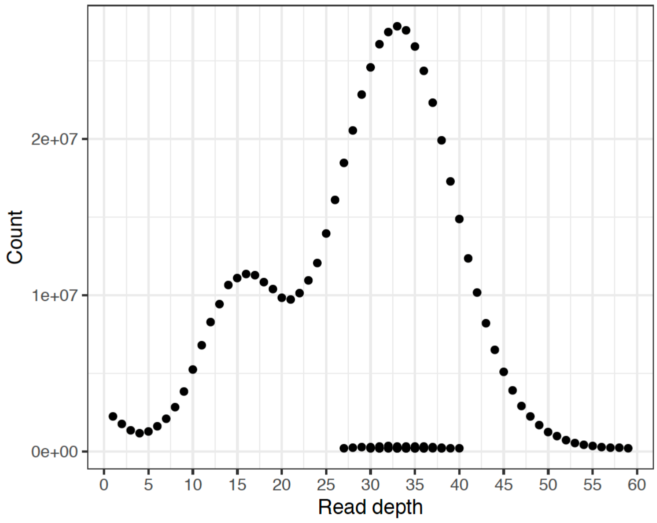


**Figure 9:** purge_haplotigs coverage histogram for *A. pisum* JIC1 Nanopore reads aligned to the Flye assembly.

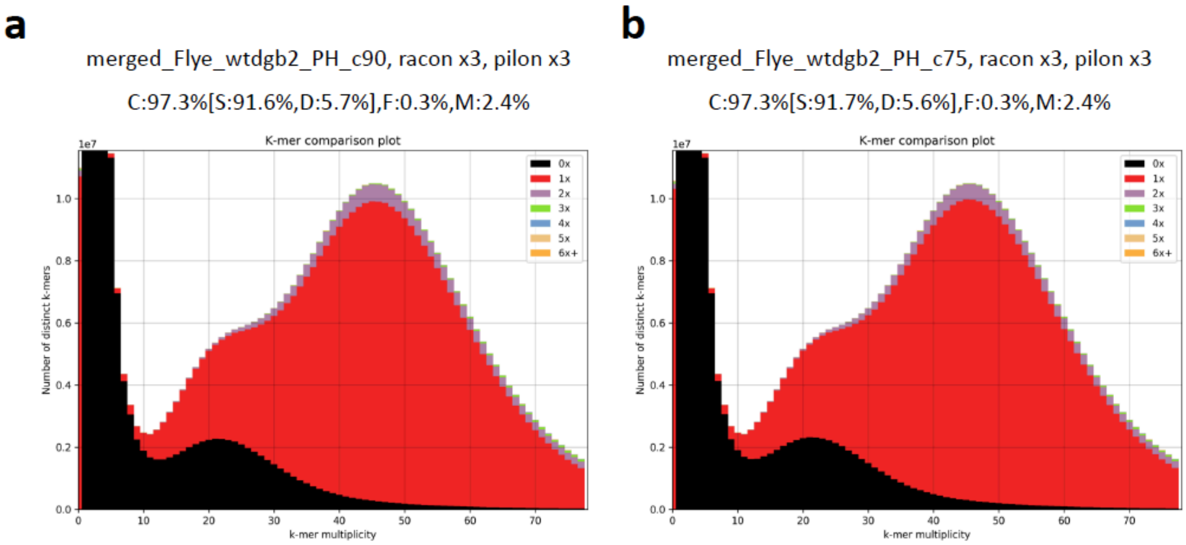


**Figure 10:** KAT k-mer spectra plots for purge_haplotigs filtered *A. pisum* JIC1 merged Flye – wtdgb2 long-read assemblies. k-mer spectra are shown for the same input assembly filtered with a 90% coverage threshold (a) or a 75% coverage threshold (**b**). BUSCO scores are shown above each of the k-mer spectra.

As we could not remove sufficient duplicated content from the merged Flye-wtdgb2 assembly, we returned to our unmerged Flye assembly and investigated whether duplicated content could be effectively filtered from this assembly. Filtering this unmerged assembly with a coverage threshold of 90% still left some residual duplicated content in the assembly (**Figure 11a**). Reducing the coverage threshold to 75% (Flye_PH_c75) more effectively removed duplicated content (**Figure 11b**) and did not result in loss of genuine single-copy content from the assembly. Using the 75% coverage threshold, ~50 Mb of content was removed by purge_haplotigs (**Table 4**), bringing it close to the predicated *A. pisum* genome size of 514 Mb (Wenger *et al.* 2017). Even though the Flye_PH_c75 assembly was less contiguous than the merged assemblies or the stand alone wtdgb2 assemblies, we selected the former assembly for downstream scaffolding, because k-mer analysis and BUSCO showed that it is the most accurate haploid representation of the genome i.e. it is complete assembly and has low levels of duplication.

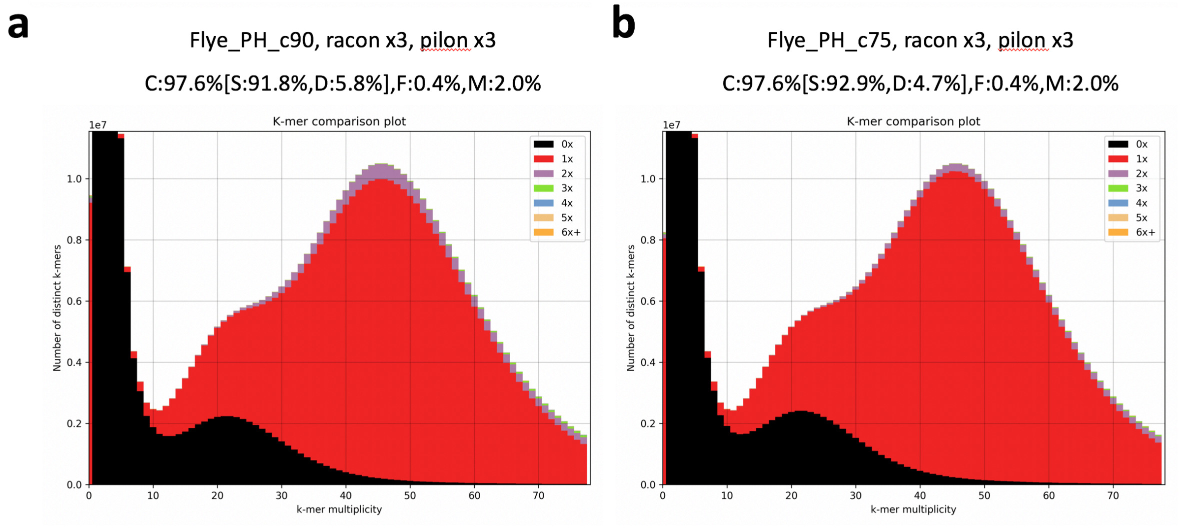


**Figure 11:** KAT k-mer spectra plots for purge_haplotigs filtered *A. pisum* JIC1 Flye long-read assemblies. k-mer spectra are shown for the same input assembly filtered with a 90% coverage threshold (**a**) or a 75% coverage threshold (**b**). BUSCO scores are shown above each of the k-mer spectra.

##### *10X genomics linked-read scaffolding*

We further improved the contiguity of the Flye_PH_c75 assembly by scaffolding it with 10X Genomics linked-reads. We applied an iterative strategy using Scaff10x v4 (https://github.com/wtsi-hpag/Scaff10X) and Tigmint v1.1.2 (Jackman *et al.* 2018). We first carried out a parameter sweep with Scaff10x to identify settings that maximised contiguity, exploring values for the "-edge" and "-block" parameters between 10 Kb and 50 Kb (**Table 5**). The most contiguous assembly was achieved with "-edge" and "-block" set to 45 Kb. With these parameters, the N50 of the Flye_PH_c75 was increased from 0.5 Mb to 2.96 Mb and the longest scaffold was 18.82 Mb. Following recommendations in the Scaff10x documentation, a second round of scaffolding was run with the same parameters, giving a marginal increase in contiguity of an N50 of 2.97 Mb. Subsequently, misassembled scaffolds were broken up using tigmint and standard settings followed by a final round of scaffolding with ARCS (Yeo *et al.* 2018). This resulted in a further increase in contiguity, raising the N50 to 3.72 Mb (Table 5). The longest scaffold in the tigmint_arcs assembly was 26.19 Mb.

**Table 5:** Scaff10x parameter optimisation and iterative scaffolding of the Flye_PH_c75 contig assembly. Green shading highlights the assemblies used in downstream steps.

| Assembly | N50 (Mb) | L50 | Longest scaffold (Mb) |
|---|---|---|---|
| scaff10x_edge_block_10000 | 0.73 | 186 | 5.79 |
| scaff10x_edge_block_15000 | 1.34 | 88 | 5.79 |
| scaff10x_edge_block_20000 | 2.13 | 65 | 11.98 |
| scaff10x_edge_block_25000 | 2.16 | 60 | 12.27 |
| scaff10x_edge_block_30000 | 2.42 | 49 | 18.82 |
| scaff10x_edge_block_35000 | 2.44 | 50 | 18.82 |
| scaff10x_edge_block_40000 | 2.44 | 52 | 19.03 |
| scaff10x_edge_block_45000 | 2.96 | 44 | 18.82 |
| scaff10x_edge_block_50000 | 2.73 | 44 | 18.82 |
| scaff10x_edge_block_45000_round_2 | 2.97 | 42 | 18.82 |
| tigmint_arcs | 3.71 | 35 | 26.19 |

##### *HiC scaffolding*

HiC scaffolding was carried out following the procedure outlined for *M. persicae* clone O above, with the exception that we modified the 3D-DNA parameter "--editor-repeat-coverage" to 4 to avoid overly aggressive breaking of the input assembly during the misassembly detection step. Manual inspection of the HiC contact map after initial scaffolding with 3D-DNA revealed only minor corrections were needed in JBAT (**Figure 12a** and **b**).

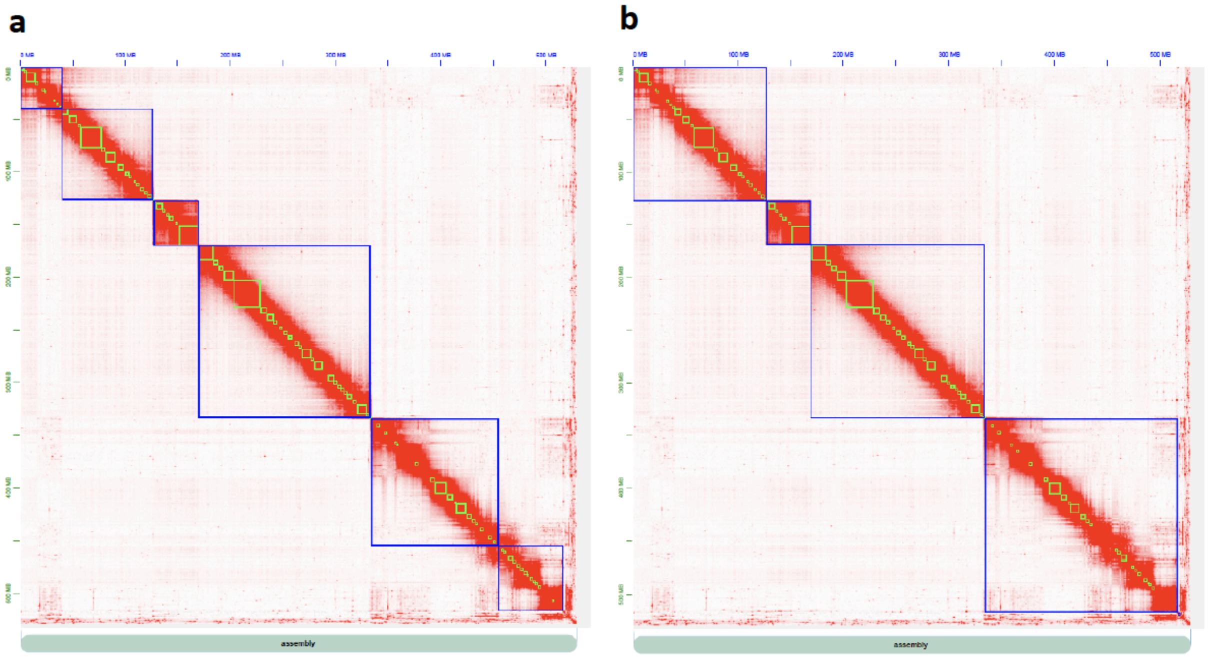

**Figure 12:** Heatmaps showing the frequency of HiC contacts along the 3d-DNA scaffolded *A. pisum* clone O 10X scaffolded contig assembly before (**a**) and after (**b**) manual review with JBAT. Blue lines indicate super scaffolds and green lines show input scaffolds from the 10x scaffolded contig assembly. The X and Y axis show cumulative length in Mb.

##### *Contamination filtering and final quality control*

Contamination filtering with BlobTools and final quality control steps for *A. pisum* JIC1 were carried out as described for *M. persicae* clone O with the exception that we used our 10X Genomics reads instead of PCR free Illumina reads to estimate scaffold coverage. Low levels of contamination were found upon manual inspection of the BlobTools taxon annotated GC content-coverage plots (**Figures 13** and **14**). A single short scaffold assigned to the obligate endosymbiont *Buchnera aphidicola* (n = 1, total content = 23 Kb; a short fragment of the genome) and 10 scaffolds assigned to the secondary symbiont *Serratia symbiotica* (n = 10, total content = 2.71 Mb) were removed from the final assembly. Other scaffolds that were removed from the assembly correspond to low coverage contamination based on the criteria of having less than 10x average coverage in the Illumina reads and the Nanopore long-reads (n = 94, total content = 1.15 Mb). Finally, to generate the final *A. pisum* clone O v2 release, the remaining scaffolds were renamed and ordered by size with SeqKit. The assembly was then checked a final time with KAT comp and BUSCO (**Supplementary Figures 2** and **3**).

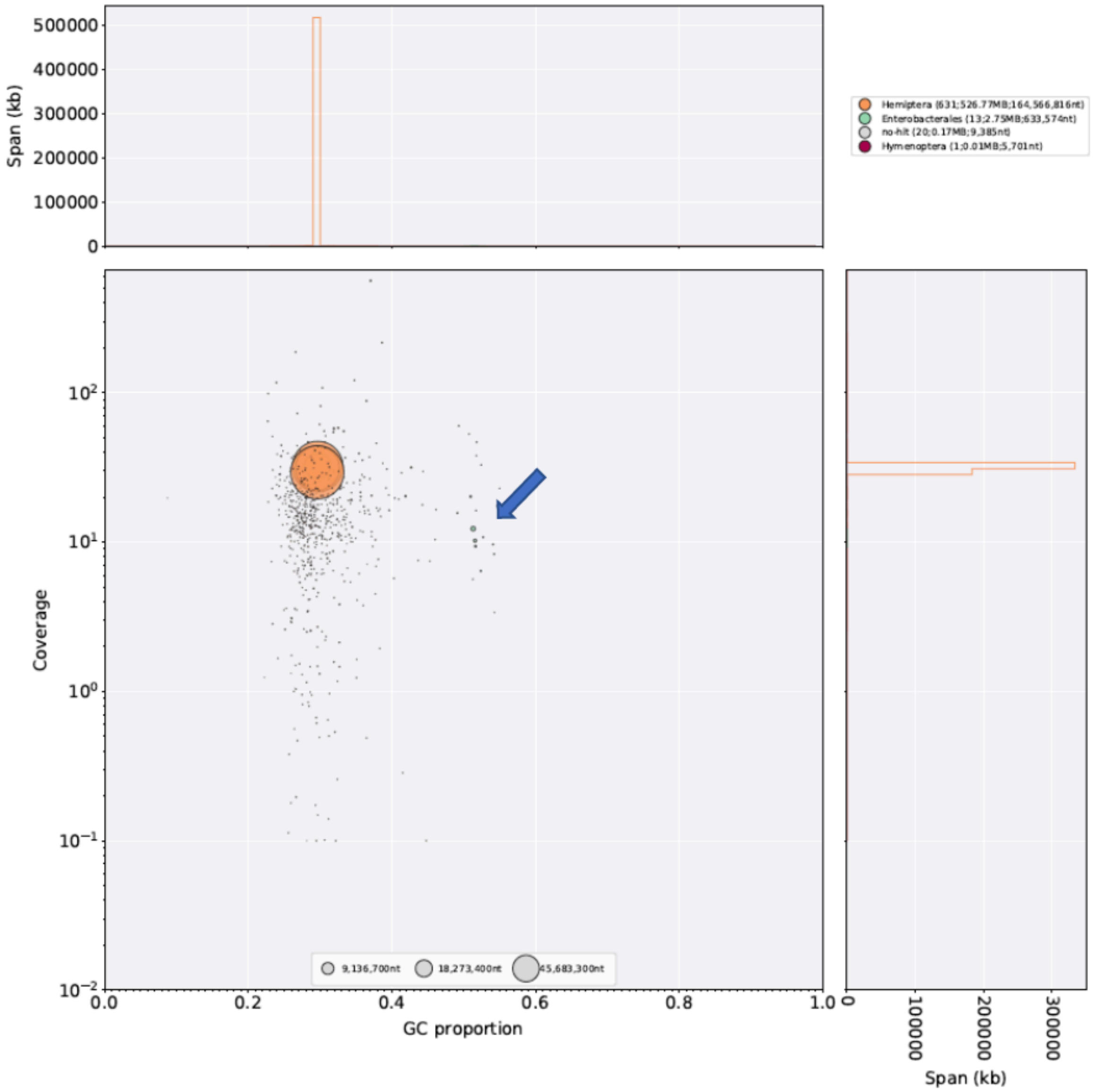


**Figure 13:** Taxon-annotated GC content-coverage plot of the *A. pisum* JIC1 HiC scaffolded assembly. Each circle represents a scaffold in the assembly, scaled by length, and coloured by order-level NCBI taxonomy assigned by BlobTools. The X axis corresponds to the average GC content of each scaffold and the Y axis corresponds to the average coverage based on alignment of Nanopore long-reads. Marginal histograms show cumulative genome content (in Kb) for bins of coverage (Y axis) and GC content (X axis). The blue arrow indicates scaffolds derived from the secondary symbiont *Serratia symbiotica*.

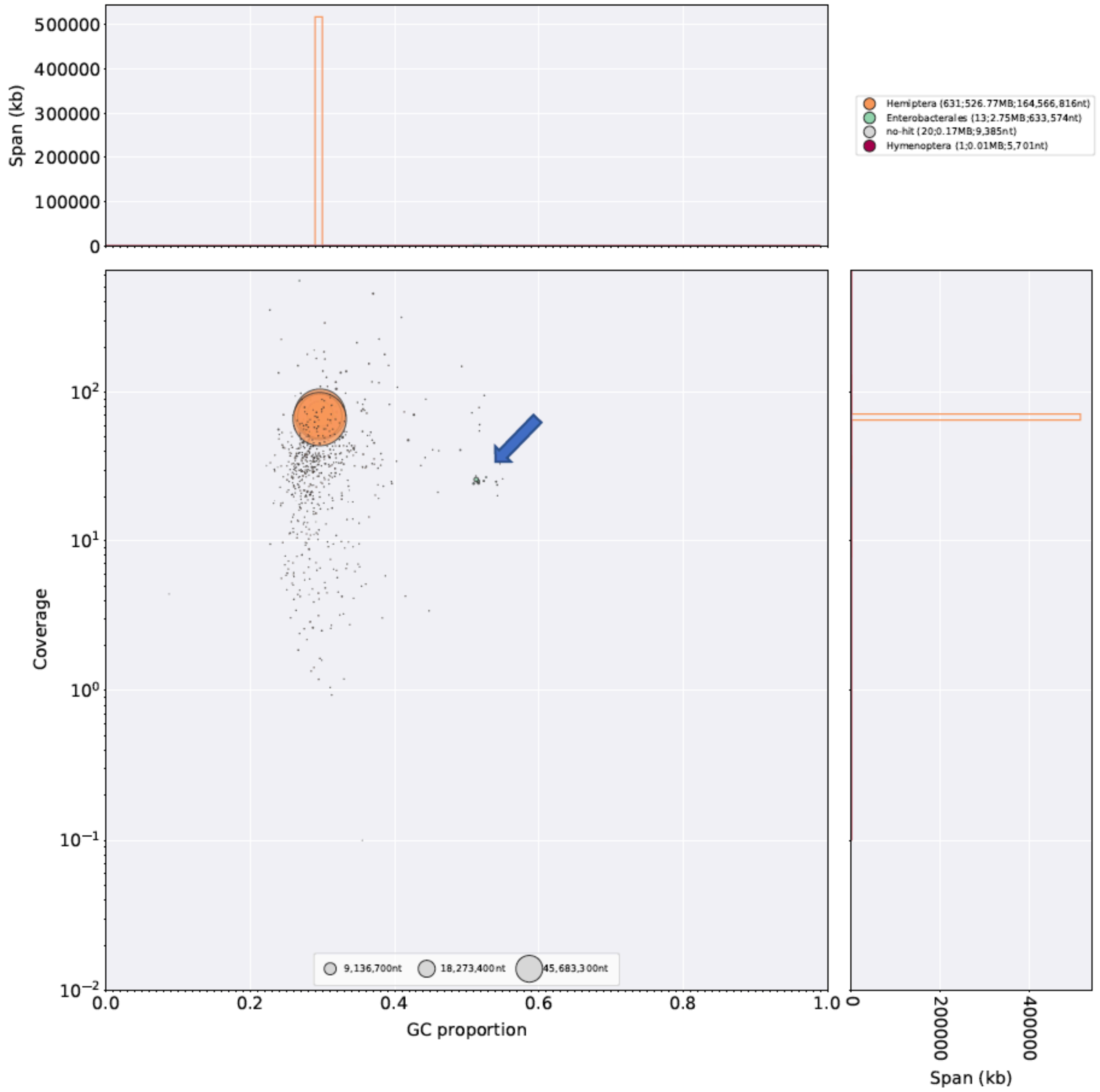


**Figure 14:** As for Figure 13 but using *A. pisum* JIC1 10X Genomics reads to calculate scaffold coverage.
